# A light-responsive lipid nanoparticle platform delivering Cas9 RNP for *VEGFA* editing suppresses choroidal neovascularisation

**DOI:** 10.64898/2026.10.24.761602

**Authors:** Rui Sang, Xinpu Yang, Ewa Goldys, Yagiz Alp Aksoy, Gerald Liew, Wei Deng

**Affiliations:** School of Biomedical Engineering, Faculty of Engineering, UNSW Sydney, NSW 2052, Australia; School of Electrical, Mechanical and Biomedical Engineering, FEIT, University of Technology Sydney, Ultimo, NSW 2007, Australia; Biomedical AI Centre, Centenary Institute and Faculty of Medicine and Health, The University of Sydney; Centre for Vision Research, Department of Ophthalmology, and Westmead Institute for Medical Research, The University of Sydney; Department of Ophthalmology, Westmead Hospital.

**Keywords:** Gene editing, Light-responsive lipid nanoparticles, Verteporfin, VEGFA, Choroidal neovascularisation

## Abstract

Choroidal neovascularisation (CNV) is a major cause of irreversible vision loss. Conventional anti-VEGF therapies for CNV often require repeated intravitreal injections, imposing substantial treatment burden and cumulative injection-related risk. Genome editing to disrupt *VEGFA* could provide durable suppression, potentially from a single administration. However, durability depends on permanently disrupting the target in a well-controlled way, as unrestricted *VEGFA* loss can cause choriocapillaris atrophy and compromise retinal function. This represents a major safety barrier to clinical translation. In this study, we developed lipid-based nanoparticles to release Cas9/sgRNA ribonucleoprotein (RNP) complexes under light illumination, enabling light-controlled editing of the *VEGFA* gene. The nanoparticles had a mean hydrodynamic diameter of approximately 130 nm with low polydispersity (0.296 ± 0.032) and high sgRNA encapsulation efficiency (90.33 ± 0.15%). In human retinal pigment epithelial cells, light exposure increased the extent of *VEGFA* disruption relative to the non-illuminated condition, as confirmed by targeted next-generation sequencing, with corresponding reductions in *VEGFA* mRNA and protein expression. Following a single intravitreal injection in a laser-induced rat CNV model, the nanoparticles combined with light illumination significantly reduced vascular leakage and CNV-associated structural abnormalities compared with both vehicle and non-illuminated controls, and ocular safety assessment revealed no apparent nanoparticle-related structural abnormalities in the retina. This study provides a foundation for spatially controlled *VEGFA* genome editing as a potential therapeutic strategy for *VEGFA*-driven macular neovascular disease.

---

Neovascular age-related macular degeneration (nAMD) is a major cause of irreversible vision loss and is characterised by pathological choroidal neovascularisation (CNV), vascular leakage and progressive disruption of retinal architecture ^1, 2^. Upregulation of vascular endothelial growth factor A (*VEGFA*) is a central driver of pathological ocular angiogenesis, and its clinical management typically involves intravitreal administration of VEGF inhibitors such as ranibizumab, aflibercept and faricimab ^3–5^. However, these anti-VEGF agents provide only transient suppression of neovascular activity, requiring frequent (e.g monthly or bimonthly) and often lifelong intravitreal injections. Repeated treatment entails substantial financial costs, with these agents consistently some of the most expensive medications in pharmacy formularies, and exposes patients to cumulative risks of injection-related complications, including elevated intraocular pressure, intraocular haemorrhage, uveitis and endophthalmitis. ^6–8^. *In vivo* CRISPR-based gene therapies offer the potential to overcome this limitation by enabling sustained therapeutic effects after one or a few administrations ^9, 10^. However, achieving these effects through permanent target disruption raises safety concerns, as unrestricted *VEGFA* loss can cause choriocapillaris atrophy and compromise retinal function ^11–13^, posing a major barrier to clinical translation.

Early studies applied adeno-associated virus (AAV) to deliver CRISPR components to retinal tissues for CNV management ^14, 15^. AAV-mediated delivery of Streptococcus pyogenes Cas9 (SpCas9) together with *VEGFA*-targeting sgRNAs achieved efficient suppression of *VEGFA* expression and pathological neovascularisation in a mouse model ^14^. Despite these promising results, viral delivery carries several intrinsic constraints, including host immunogenicity and off-target editing arising from prolonged intracellular expression of the genome-editing ^16–18^. Most importantly, constitutive promoters generally lack disease- or lesion-specific control over transgene expression ^19–21^. Nonviral lipid-based nanoparticles provide an alternative platform for gene delivery, with tunable physicochemical properties, capacity for diverse genome-editing cargos and favourable biocompatibility ^22–24^. Recent advances have enabled CRISPR delivery to ocular tissues, including Cas9 mRNA and sgRNA for *VEGFA* editing ^25–29^. However, existing lipid nanoparticles lack an intrinsic mechanism for selectively delivering their cargo to diseased retinal lesions while sparing healthy tissue. This is particularly challenging because the same cell types are present in both regions, and reliable lesion-specific biomarkers for selective targeting are limited. This highlights the need for a delivery system that enables spatially restricted cargo release and confines therapeutic activity to the targeted lesion.

Stimuli-responsive nanocarriers provide a potential strategy by enabling cargo release in response to defined biological or external triggers ^30, 31^. For ocular applications, visible light is particularly attractive because the eye is readily accessible to external illumination, allowing cargo release to be spatially restricted to the illuminated area ^32–34^. In our previous studies, we developed a light-triggered liposomal delivery system in which verteporfin (VP), a clinically used photosensitiser, generated reactive oxygen species (ROS) upon light activation, triggering disruption of the liposomal membrane and intracellular cargo release ^35–37^. Incorporating Cas9 RNP targeting *VEGFA* into this light-triggered liposomal system could enable spatially precise gene editing in the retina by confining Cas9 RNP activity to the illuminated area while sparing surrounding non-illuminated retinal tissue.

In this study, we built on this platform by developing a Cas9/sgRNA-loaded liposomal formulation (Lipo-Cas9-VP) targeting *VEGFA* and evaluating its light-controlled genome-editing activity in retinal cells *in vitro* and *in vivo*. We systematically characterised the formulation and evaluated its cellular uptake, biocompatibility and light-dependent genome-editing activity in human retinal pigment epithelial cells. We then assessed the *in vivo* ocular safety of intravitreal administration of this formulation, followed by a proof-of-concept therapeutic evaluation in a laser-induced CNV rat model using fluorescein angiography and optical coherence tomography. This study provides a strategy for developing spatially precise gene therapy for *VEGFA*-driven macular diseases by integrating light-responsive Cas9 RNP delivery with external light control.

**Scheme 1.**
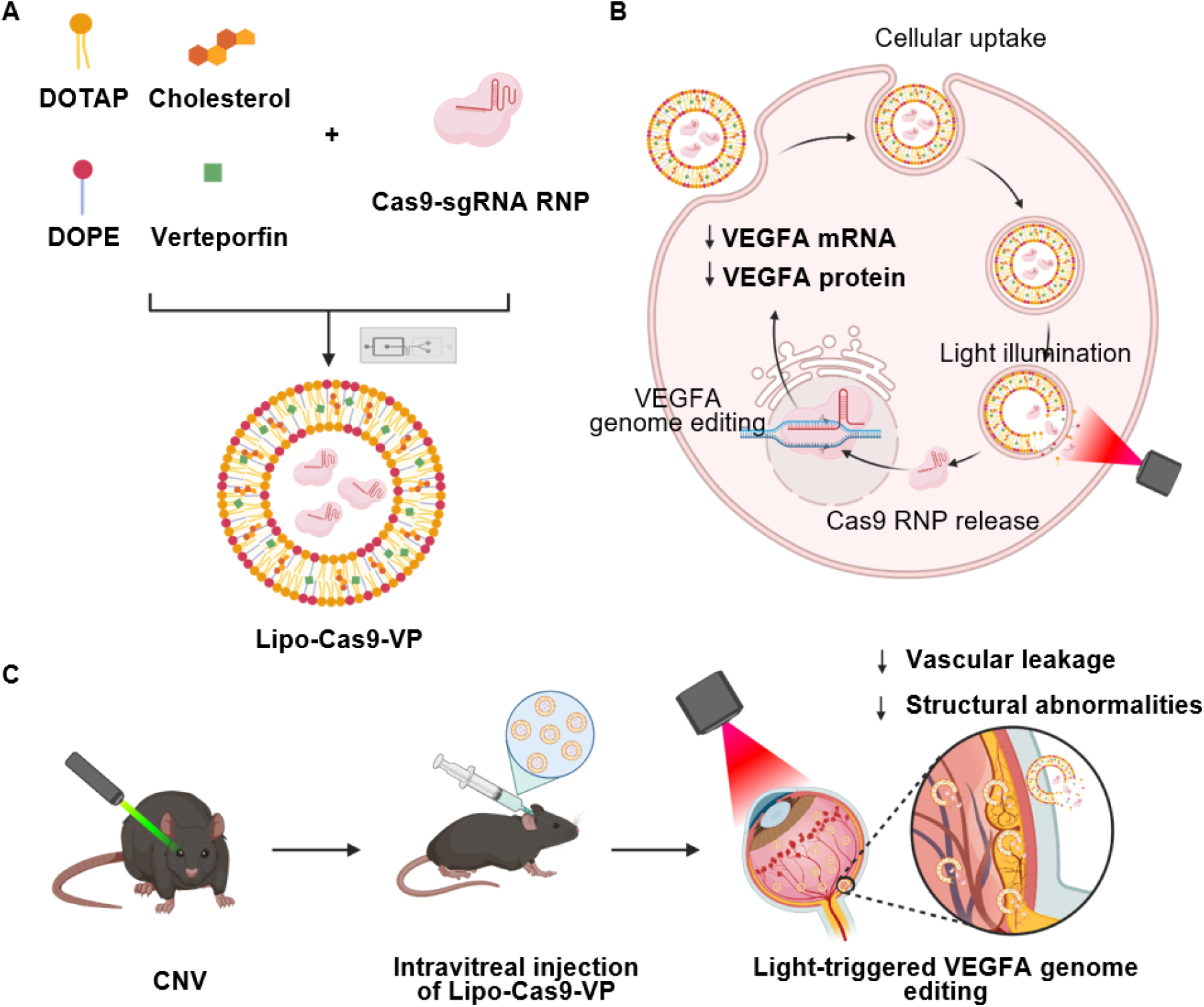
Schematic illustration of light-responsive liposome platform (Lipo-Cas9-VP) for CNV treatment. (A) Preparation of Lipo-Cas9-VP, (B) Light-controlled cargo release and activity *in vitro* and (C) Light-triggered *VEGFA* genome editing in a rat model of laser-induced CNV.

## Results

### Physicochemical characterisation of Lipo-VP and Lipo-Cas9-VP

Cryogenic transmission electron microscopy (cryo-TEM) revealed that Lipo-VP predominantly exhibited a hollow spherical morphology with particle dimensions of approximately 100 nm (Figure 1A). Lipo-VP exhibited a mean hydrodynamic diameter of 121.4 ± 0.87 nm and a PDI of 0.165 ± 0.032. Incorporation of Cas9/sgRNA RNP increased the mean hydrodynamic diameter to 132.3 ± 1.04 nm and PDI to 0.296 ± 0.032 (Table 1). Both formulations exhibited a positive surface charge, although RNP loading reduced the zeta potential from +34.50 ± 1.33 mV for Lipo-VP to +21.10 ± 1.04 mV for Lipo-Cas9-VP (Table 1). Cas9 protein was pre-complexed with sgRNA to form the RNP prior to liposome formulation. The sgRNA encapsulation efficiency (EE%) was 90.33 ± 0.15%, as determined using a RiboGreen RNase protection assay, providing an indication of efficient encapsulation of the pre-formed RNP within the liposomal formulation.

**Figure 1.**
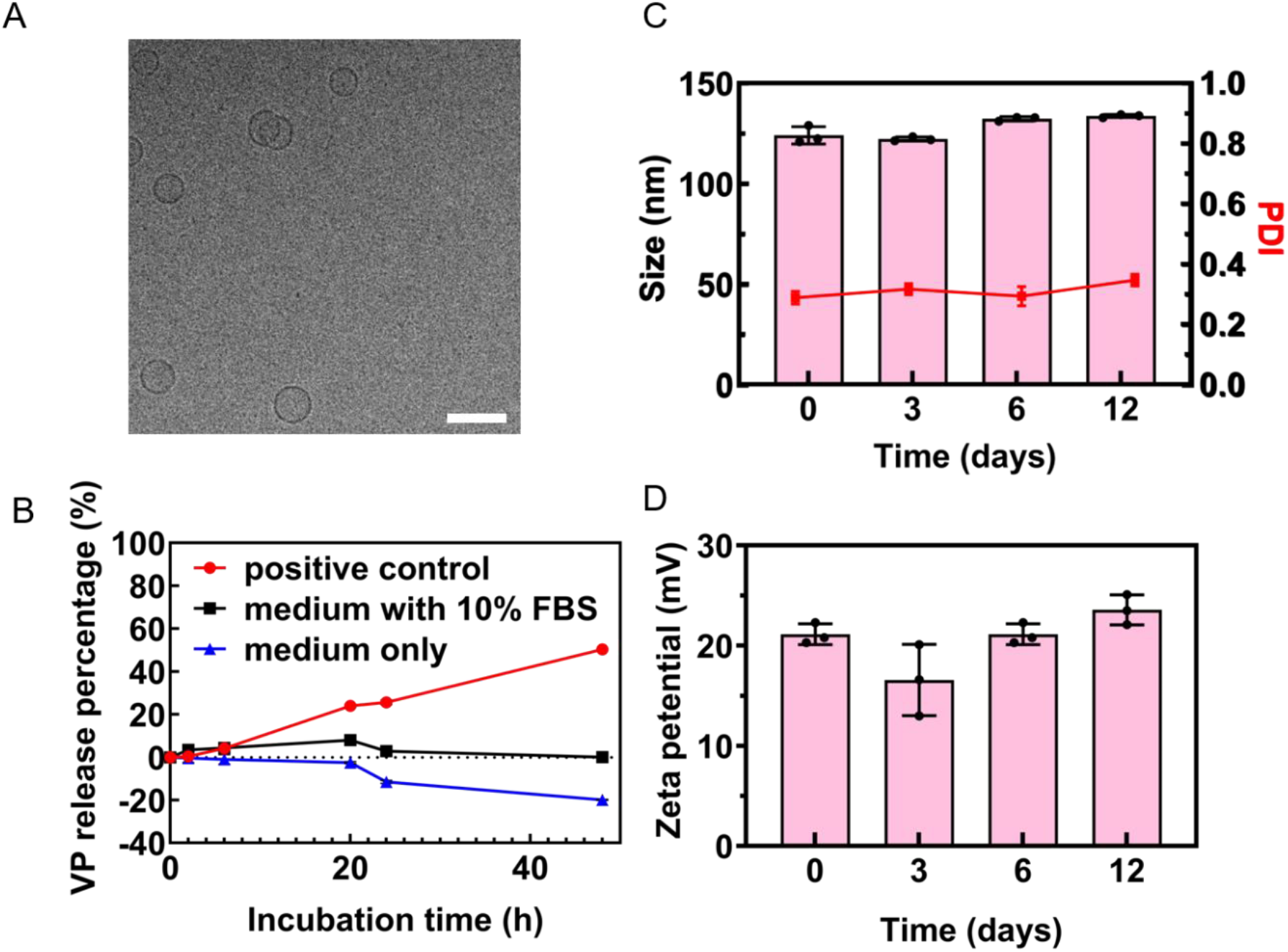
(A) Representative Cryo-TEM image of Lipo-VP, Scale bar is 200 nm; (B) VP release from Lipo-VP in the presence or absence of FBS over 48 h; (C&D) Colloidal stability of Lipo-Cas9-VP in Tris buffer over 12 days at 4 °C, assessed by changes in (C) hydrodynamic diameter and PDI and (D) zeta potential. Data are presented as mean ± SD (n = 3).

**Table 1.**
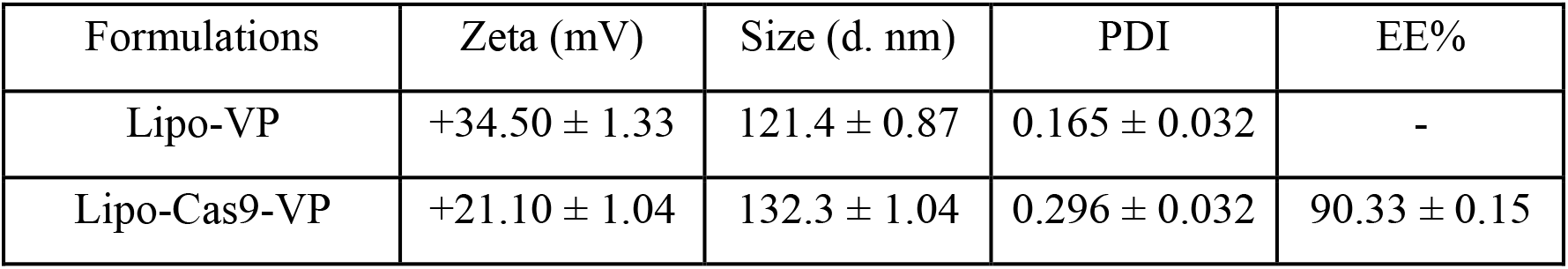
Physicochemical characteristics of Lipo-VP and Lipo-Cas9-VP formulations.

| Formulations | Zeta (mV) | Size (d. nm) | PDI | EE% |
| --- | --- | --- | --- | --- |
| Lipo-VP | $+34.50 \pm 1.33$ | $121.4 \pm 0.87$ | $0.165 \pm 0.032$ | - |
| Lipo-Cas9-VP | $+21.10 \pm 1.04$ | $132.3 \pm 1.04$ | $0.296 \pm 0.032$ | $90.33 \pm 0.15$ |

The stability of the formulations was subsequently evaluated under biologically relevant and storage conditions. VP retention was assessed by dialysis in the presence or absence of FBS over 48 h. Minimal VP release was observed throughout the incubation period under both conditions (Figure 1B), indicating that the nanoparticles remained stable in the presence of serum. The colloidal stability of Lipo-Cas9-VP was further assessed by monitoring particle size, PDI and zeta potential over 12 days. No significant changes in these parameters were observed during the study period (Figure 1C and 1D), indicating that Lipo-Cas9-VP remained physicochemically stable during storage in Tris buffer at 4 °C for at least 12 days, without detectable particle aggregation.

### Cellular uptake and cell viability of Lipo-VP in ARPE19 cells

Cellular uptake of Lipo-VP by ARPE19 cells was evaluated using confocal fluorescence microscopy. As shown in Figure S1, intracellular VP fluorescence increased progressively with incubation time, indicating time-dependent accumulation of Lipo-VP in ARPE19 cells. Fluorescence intensity approached a plateau after approximately 3 h, and this incubation time was therefore selected for subsequent experiments. The effect of Lipo-VP concentration on cellular uptake was then evaluated. Increasing the formulation concentration resulted in greater intracellular VP fluorescence, with the 221.44 µg/mL of Lipo-VP producing a much stronger fluorescence signal than the 110.72 µg/mL of Lipo-VP (Figure 2).

**Figure 2.**
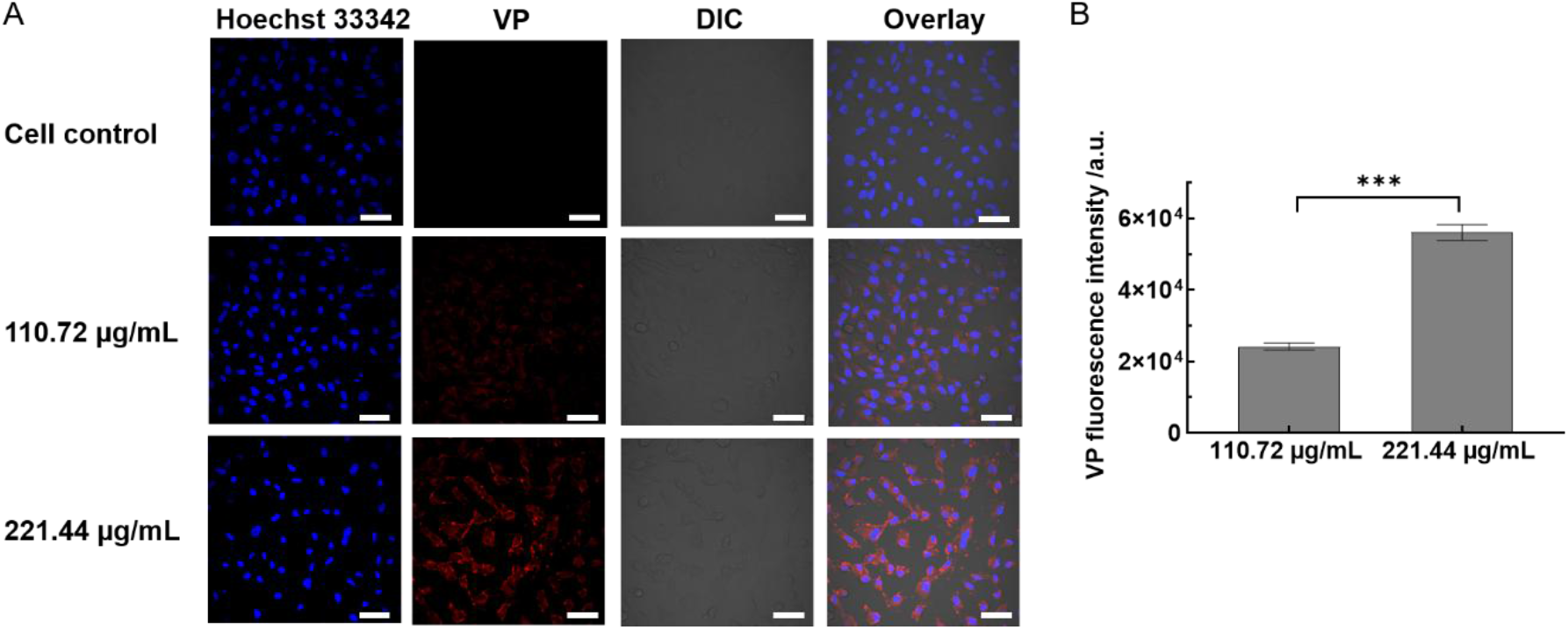
Cellular uptake of Lipo-VP by ARPE19 cells. (A) Representative confocal fluorescence images of ARPE19 cells incubated with different concentration of Lipo-VP (110.72 µg/mL and 221.44 µg/mL) for 3 hr. Scale bar is 50 μm; (B) Quantitative analysis of intracellular VP fluorescence in ARPE19 cells at different incubation concentrations as shown in Figure 2A; Mean ± SD (n=3), *p < 0.05, **p < 0.01 and ***p < 0.005.

Cell viability following Lipo-VP treatment was subsequently assessed using an MTS assay. In the absence of light irradiation, treatment at concentrations of 55.36, 110.72 and 221.44 µg/mL did not significantly affect ARPE19 cell viability (Figure S2). The effects of Lipo-VP and light irradiation were then evaluated in combination. Light irradiation alone did not significantly affect cell viability at any of the tested exposure times (Figure S3). Similarly, cells treated with the 110.72 µg/mL of Lipo-VP maintained greater than 90% viability following light irradiation for up to 8 min. At 221.44 µg/mL of Lipo-VP, cell viability remained high following 4 min of irradiation (92.00 ± 7.47%), while a longer irradiation period resulted in a further reduction in viability. Based on the efficient cellular uptake at a Lipo-VP concentration of 221.44 µg/mL and the high cell viability maintained following 4 min of light irradiation, these conditions were selected for subsequent *in vitro* experiments.

### Selection and validation of a *VEGFA*-targeting sgRNA

Before encapsulation of Cas9-sgRNA RNP complexes into Lipo-VP, four candidate sgRNAs targeting human *VEGFA* were screened in ARPE19 cells. To enable reliable assessment of *VEGFA* expression and genome-editing efficiency, the primer pairs for RT-qPCR and T7E1 analysis were first optimised. Six candidate RT-qPCR primer pairs were evaluated for amplification efficiency and specificity using melting-curve analysis, agarose gel electrophoresis and Sanger sequencing. Primer pair 3 was selected based on its consistent amplification performance and confirmed target specificity (Table S6 and Figure S4). Similarly, four primer pairs targeting the genomic region encompassing the *VEGFA* target site were evaluated, and primer pair 3 was selected for subsequent T7E1 analysis (Figure S5).

The four candidate sgRNAs were subsequently evaluated 72 h after transfection using the optimized assays. RT-qPCR analysis showed that sgRNA002 significantly reduced *VEGFA* mRNA expression compared with untreated cells (Figure 3A). Consistent with the RT-qPCR results, T7E1 analysis indicated that sgRNA002 produced the highest genome-editing efficiency (53.89 ± 1.18%), compared with sgRNA001 (47.27 ± 1.52%), sgRNA003 (21.54 ± 5.46%) and sgRNA004 (19.13 ± 4.11%). The untreated control exhibited a background signal corresponding to an apparent editing efficiency of 10.30 ± 3.80% (Figure 3B and Figure S6). The predicted specificity of sgRNA002 was further assessed using the genome-wide Cas-OFFinder tool (http://www.rgenome.net/cas-offinder/) ^38^. Potential off-target sites were screened by allowing up to three mismatches to the intended target sequence and requiring an adjacent NGG protospacer-adjacent motif (PAM) ^39^. No potential off-target sites meeting these criteria were identified (Table S5). Based on its efficient *VEGFA* mRNA suppression, high estimated genome-editing activity and favourable predicted specificity, sgRNA002 was selected for formation of the Cas9/sgRNA RNP complex and subsequent encapsulation into the Lipo-VP formulation for evaluation of light-triggered genome editing.

**Figure 3.**
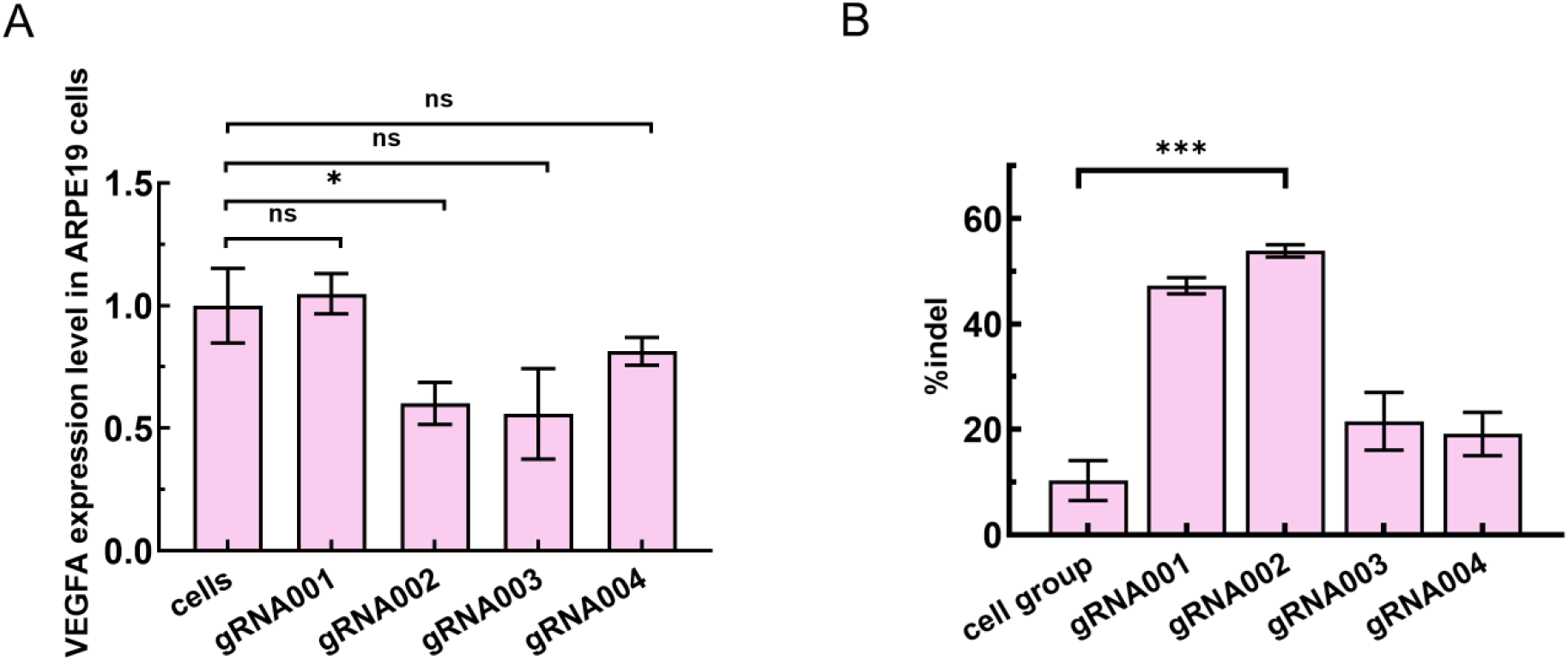
Screening and selection of sgRNAs targeting *VEGFA* in ARPE19 cells. (A) Relative *VEGFA* mRNA expression and (B) estimated genome-editing efficiency determined by T7E1 assay in ARPE19 cells 72 h after transfection with Cas9/sgRNA RNPs containing different sgRNAs using Lipofectamine CRISPRMAX. Data are presented as mean ± SEM (A) or mean ± SD (B) (n = 3), *p < 0.05, **p < 0.01 and ***p < 0.005.

### Light-triggered genome editing of *VEGFA* by Lipo-Cas9-VP in ARPE19 cells

To evaluate the light-controlled genome-editing activity of Lipo-Cas9-VP, ARPE19 cells were incubated with the formulation for 3 h, followed by 4 min of light irradiation. Genome editing at the *VEGFA* locus was first assessed using the T7E1 assay. PCR amplification of the target region generated a 424-bp product, with the expected 297- and 127-bp cleavage fragments detected following T7E1 digestion (Figure S7). A relatively high apparent cleavage signal was observed in the untreated control (21.50 ± 0.76%) and in cells treated with Lipo-Cas9-VP alone (21.37 ± 1.37%). Such background signals have been reported for T7E1-based mismatch-cleavage assays and may arise from sequence variation or other assay-related factors ^40, 41^. Following Lipo-Cas9-VP treatment with light irradiation, the apparent cleavage value increased significantly to 42.21 ± 1.24% (P < 0.05; Figure 4A), consistent with light-controlled genome editing at the *VEGFA* locus.

**Figure 4.**
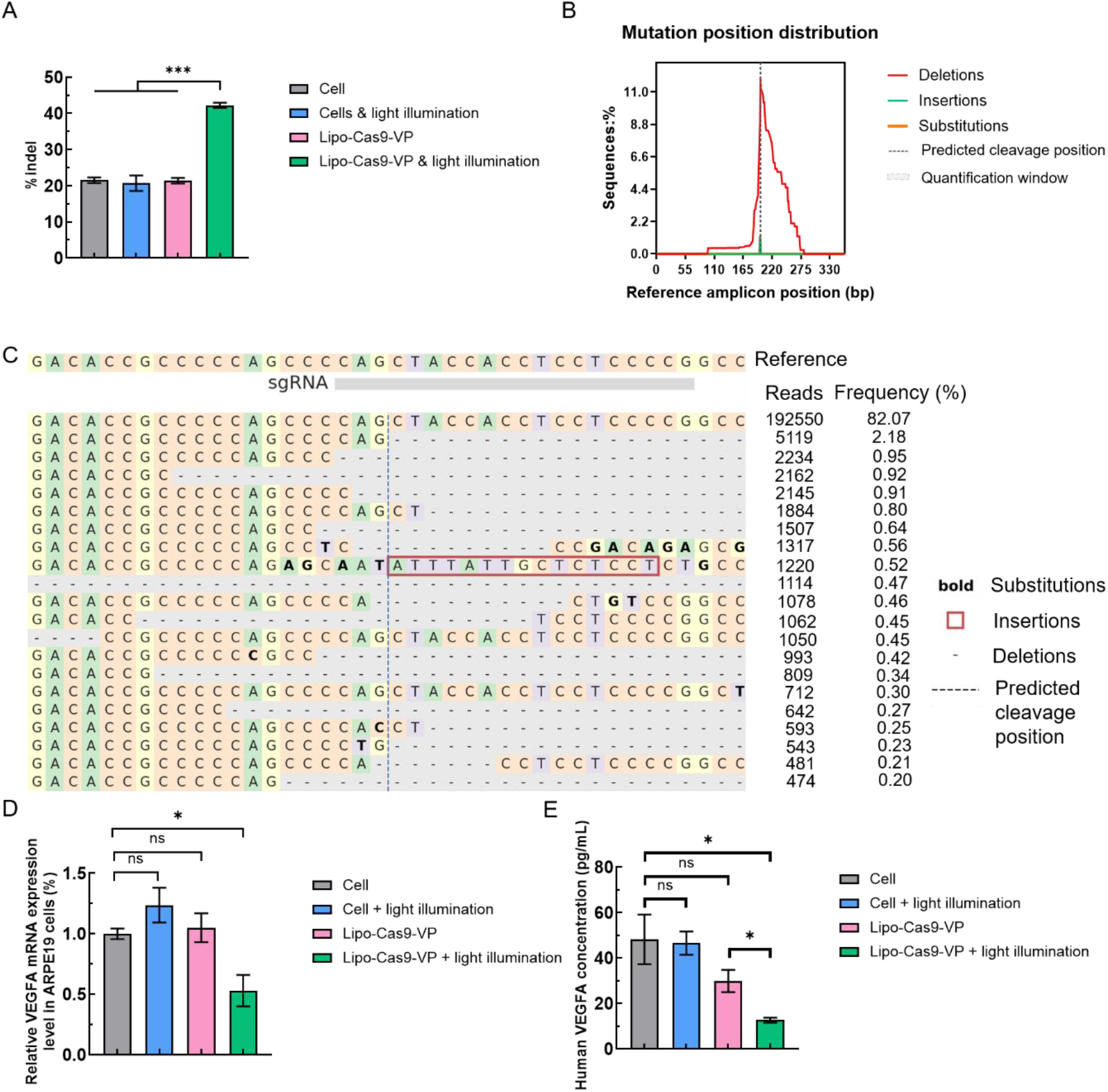
Light-triggered *VEGFA* genome editing and suppression of *VEGFA* expression in ARPE19 cells at 48 hr after treatments. (A) Estimated editing efficiency at the *VEGFA* target locus determined by T7E1 assay. (B) Position-wise distribution of insertions, deletions and substitutions following Lipo-Cas9-VP treatment with light irradiation, determined by targeted amplicon NGS. The black dashed line indicates the predicted Cas9 cleavage site, the grey dashed box the CRISPResso2 quantification window, and the grey bar the sgRNA target region. (C) Representative edited *VEGFA* alleles identified by targeted amplicon sequencing in the Lipo-Cas9-VP plus light group. Alleles with frequencies ≥ 0.2% are shown; dashes indicate deletions, red boxes insertions, bold letters substitutions and the vertical dashed line the predicted Cas9 cleavage site. (D) *VEGFA* mRNA expression measured by RT-qPCR following the indicated treatments. (E) *VEGFA* protein levels following the indicated treatments. Data are presented as mean ± SEM (D) or mean ± SD (A and E) (n = 3). *p < 0.05, ** p < 0.01, *** p < 0.005.

Targeted amplicon next-generation sequencing (NGS) was subsequently performed to further characterise editing at the *VEGFA* target locus. Sequence modifications were concentrated around the predicted Cas9 cleavage site and consisted predominantly of deletions, with insertions and substitutions detected at much lower frequencies (Figure 4B and C). Only low levels of modification (0.33%) were detected in cells treated with Lipo-Cas9-VP without light irradiation, whereas Lipo-Cas9-VP treatment followed by light irradiation resulted in an editing frequency of 13.67% (Figure S8). Together, these findings demonstrate the feasibility of light-controlled genome editing, with minimal editing in the absence of light and readily detectable editing following light activation.

The transcriptional effect of genome editing was then assessed by RT-qPCR 48 h after treatment. *VEGFA* mRNA expression was significantly reduced in cells treated with Lipo-Cas9-VP followed by light irradiation compared with both nanoparticle-only and untreated cells (Figure 4D), indicating a light-dependent suppression of *VEGFA* transcription. The effect on VEGFA protein expression was subsequently evaluated by ELISA. VEGFA protein levels were significantly lower following Lipo-Cas9-VP treatment with light irradiation than in both untreated cells and cells treated with Lipo-Cas9-VP alone (P < 0.05; Figure 4E and Figure S9). Relative to the untreated control, VEGFA protein levels were reduced from 48.18 ± 10.94 pg/mL to 12.60 ± 1.10 pg/mL following light irradiation. Together, these findings show that light-triggered genome editing at the *VEGFA* locus was accompanied by suppression of both *VEGFA* mRNA and VEGF protein expression in ARPE19 cells.

### Ocular safety of intravitreal Lipo-VP

An *in vivo* ocular safety study was conducted to evaluate the tolerability of Lipo-VP following a single bilateral intravitreal administration in a rat model (Figure 5A). No nanoparticle-related clinical abnormalities or changes in body weight were observed during the 14-day observation period (Figure S10). General ophthalmic examinations similarly showed no apparent abnormalities in either vehicle- or Lipo-VP-treated eyes throughout the study (Table S7).

**Figure 5.**
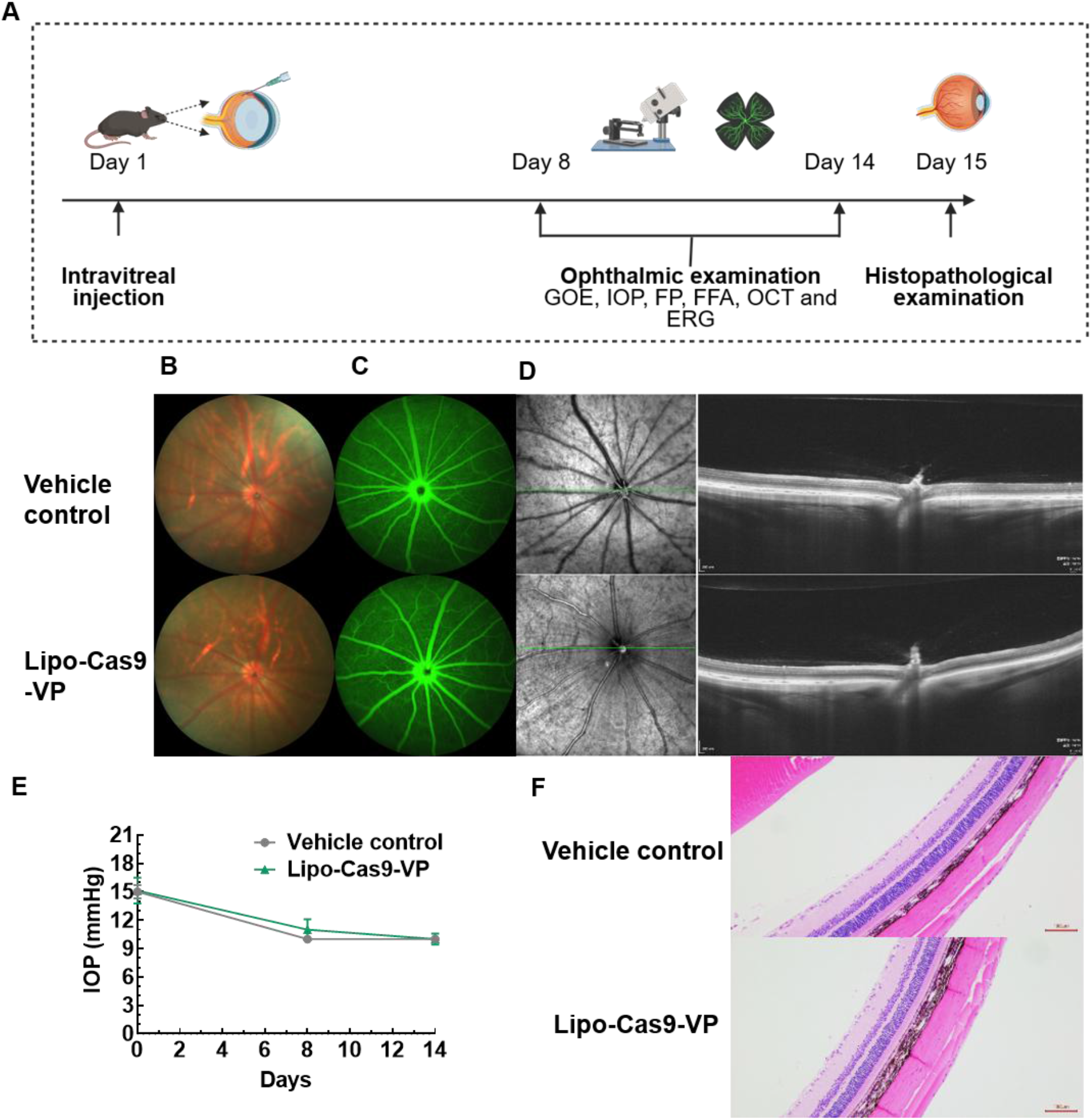
*In vivo* ocular safety of intravitreal Lipo-VP. (A) Schematic of the ocular safety study. (B-D) Representative FP images showing retinal appearance (B), FFA images (C), and OCT images (D) of vehicle- and Lipo-VP-treated eyes on Day 14. Green lines indicate the OCT scanning plane through the optic disc. (E) IOP measured at baseline (Day 0) and on Days 8 and 14 following intravitreal administration. (F) Representative H&E-stained sections of ocular tissues collected from vehicle- and Lipo-VP-treated eyes on Day 15, scale bar 100 µm. Data are presented as mean ± SD (vehicle, n = 2 rats; Lipo-VP, n = 4 rats).

Serial ophthalmic imaging revealed no apparent treatment-related ocular abnormalities following nanoparticle administration. Fundus photography showed clear ocular media and preserved optic disc and retinal vascular morphology, with no apparent differences between Lipo-VP- and vehicle-treated eyes (Figure 5B). FFA similarly showed no abnormal vascular leakage or other treatment-related angiographic changes (Figure 5C). OCT imaging demonstrated preserved retinal architecture without apparent structural abnormalities following treatment (Figure 5D). IOP remained comparable between the vehicle and Lipo-VP groups throughout the observation period, with no treatment-related increase observed on Days 8 or 14 (Figure 5E). Histopathological examination of ocular tissues collected on Day 15 also revealed no apparent treatment-related pathological changes in the eyes or optic nerves of nanoparticle-treated animals (Figure 5F).

Retinal function was further assessed by dark-adapted ERG (Table S8). On Day 8, substantial reductions in ERG amplitudes relative to baseline were observed in both the vehicle and Lipo-VP groups. For example, the DA3.0 b-wave amplitude decreased from approximately 507.88±104.28 to 185.38±68.31 μV in the vehicle group and from 490.54±109.6 to 329.93±103 μV in the Lipo-VP group. The reductions observed in both groups suggest that procedural or anaesthesia-related factors may have contributed to these changes. By Day 14, ERG amplitudes in the vehicle group showed partial recovery, whereas the DA3.0 and DA10.0 a- and b-wave amplitudes remained significantly lower in the Lipo-VP group than in the vehicle group (P ≤ 0.05). Nevertheless, these functional changes were not accompanied by corresponding abnormalities in fundus appearance, vascular leakage, retinal structure, intraocular pressure or ocular histopathology, as described above. Overall, a single intravitreal administration of Lipo-VP at 1 μg/eye was generally well tolerated over the 14-day observation period, with no consistent treatment-related adverse ocular findings across clinical observations, ophthalmic examinations, multimodal retinal imaging, IOP and histopathological assessments. However, the reductions in ERG amplitudes observed on Day 14 request further investigation in larger studies.

### Establishment of the laser-induced CNV rat model

To establish an *in vivo* model for evaluating the therapeutic efficacy of light-triggered Lipo-Cas9-VP, CNV was induced in Brown Norway rats by laser photocoagulation (Figure 6A) ^42^. Successful laser injury was confirmed by multimodal retinal imaging, including FP, FFA and OCT. Compared with healthy eyes, fundus imaging of laser-treated eyes showed clearly identifiable laser lesions surrounding the optic disc (Figure 6B). FFA further demonstrated prominent fluorescein leakage at the laser-induced lesions, consistent with disruption of the choroidal-retinal vascular barrier and development of CNV (Figure 6C). OCT further revealed focal SHRM and disruption and elevation of the outer retinal-RPE complex at the lesion sites, whereas retinal architecture remained intact in healthy eyes (Figure 6D). Collectively, these multimodal imaging findings confirmed successful establishment of the laser-induced CNV model for subsequent treatment evaluation.

**Figure 6.**
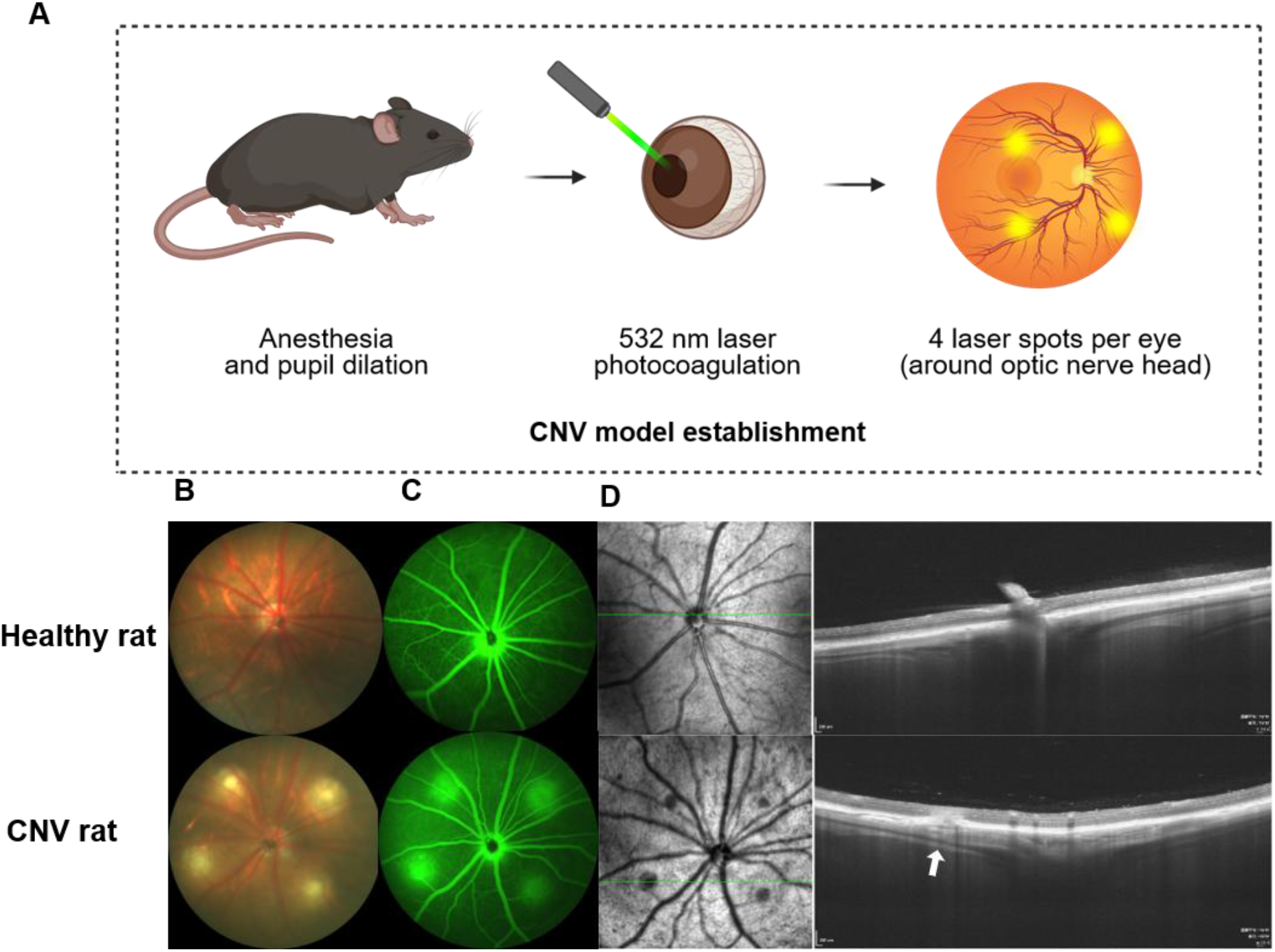
Establishment and multimodal imaging characterisation of laser-induced CNV in Brown Norway rats. (A) Schematic illustration of laser-induced CNV model establishment; (B-D) Representative FP showing retinal appearance (B), FFA images (C) and OCT images (D) of healthy rat and laser-induced CNV rat eyes (Green lines indicate the positions of the corresponding OCT B-scans, and white arrows indicate CNV-associated SHRM lesions).

Following confirmation of light-controlled genome-editing activity *in vitro*, we next investigated whether this light-controlled editing could translate into a therapeutic effect in the laser-induced CNV model. Animals received a single intravitreal administration of Lipo-Cas9-VP (1 μg/eye), with or without light irradiation, and CNV progression was assessed on Days 4 and 8 (Figure 7A). FFA was used to evaluate CNV-associated vascular leakage, with each laser-induced lesion scored according to leakage area relative to the original laser spot: Grade 0, no leakage; Grade 1, mild leakage (1-50%); Grade 2, moderate leakage (50-100%); and Grade 3, severe leakage extending beyond the laser spot (Figure 7B). On Day 4, mean leakage scores were comparable across the treatment groups. By Day 8, mean scores were 1.90 ± 0.37 for vehicle, 1.72 ± 0.33 for Lipo-Cas9-VP without light and 1.25 ± 0.27 for Lipo-Cas9-VP with light irradiation. Without light, Lipo-Cas9-VP did not significantly reduce leakage compared with vehicle (P = 0.404). In contrast, light irradiation significantly reduced leakage scores compared with both vehicle (P = 0.025) and Lipo-Cas9-VP without light (P = 0.048). These findings demonstrate a light-dependent reduction in CNV-associated vascular leakage, supporting the therapeutic potential of the spatially controlled gene editing.

**Figure 7.**
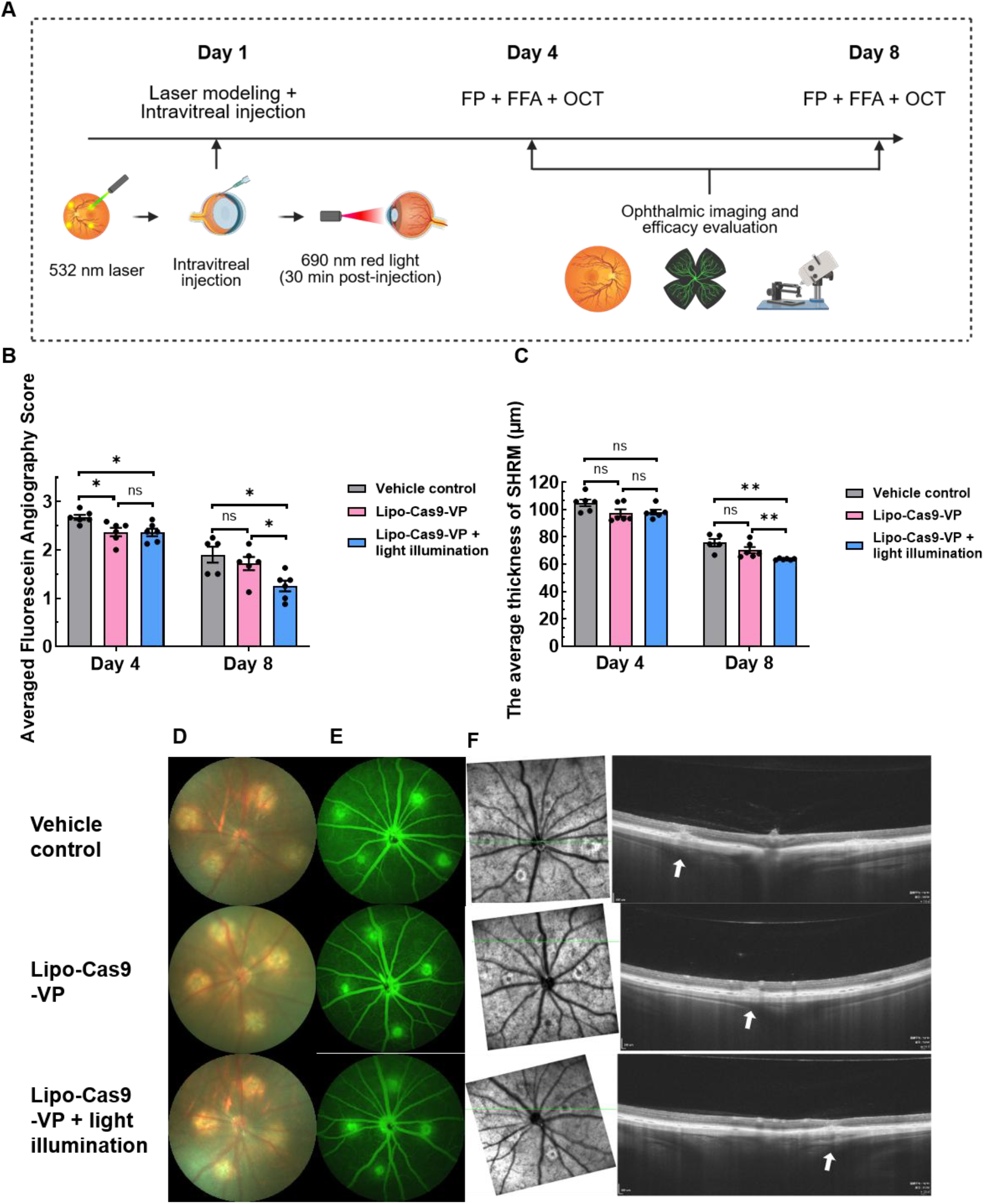
*In vivo* therapeutic effects in a laser-induced CNV model. (A) Schematic illustration of the experimental design. (B) Quantification of FFA leakage, expressed as the mean fluorescein angiography score at the indicated time points after CNV induction. (C) Quantification of CNV-associated SHRM thickness measured by OCT. (D-F) Representative FP images showing retinal appearance (D), FFA images (E) and OCT images (F) from the vehicle, Lipo-Cas9-VP and Lipo-Cas9-VP + light groups on Day 8. Green lines indicate the OCT scanning plane through the optic disc. White arrows indicate CNV-associated SHRM lesions on OCT. Data are presented as mean ± SEM (n = 6 rats).

A consistent light-dependent effect was observed by OCT (Figure 7C). Mean SHRM thickness was comparable among groups on Day 4. By Day 8, SHRM thickness was 75.81 ± 6.08 μm, 70.36 ± 5.66 μm and 63.76 ± 0.58 μm in the vehicle, Lipo-Cas9-VP and Lipo-Cas9-VP + light groups, respectively. The Lipo-Cas9-VP + light group showed significantly lower SHRM thickness than both the vehicle (P = 0.009) and Lipo-Cas9-VP groups (P = 0.006), indicating a significant reduction in CNV-associated pathology at Day 8.

Consistent with the quantitative analyses, representative images obtained on Day 4 (Figure S11) and Day 8 showed prominent laser-induced lesions, fluorescein leakage and focal SHRM in vehicle-treated eyes (Figure 7D-F). No treatment-related clinical abnormalities or adverse effects on body weight were observed during the 8-day observation period (Figure S12). Together, the FFA and OCT findings indicate that light-dependent gene editing enhanced the therapeutic effects, resulting in reduced vascular leakage and CNV-associated structural pathology.

### Discussion and conclusion

In this study, we developed light-responsive liposomes loaded with Cas9/sgRNA RNPs targeting *VEGFA*, providing initial evidence for a light-controlled genome-editing approach for the treatment of CNV. The resulting Lipo-Cas9-VP formulation was readily internalized by ARPE19 cells. Importantly, *VEGFA* editing showed a clear dependence on light exposure. Targeted amplicon NGS detected an editing frequency of 13.67% following Lipo-Cas9-VP treatment with light, compared with only 0.33% in the absence of light, with deletions predominantly concentrated around the predicted Cas9 cleavage site. This increase in genome editing was accompanied by reductions in *VEGFA* mRNA and protein expression *in vitro*. The therapeutic potential of this strategy was further assessed in a laser-induced CNV rat model. Light irradiation of Lipo-Cas9-VP-treated eyes resulted in significant reductions in both CNV-associated vascular leakage and structural pathology, supporting the therapeutic potential of the light-controlled approach.

The degree of *VEGFA* disruption achieved here was moderate rather than complete, and this is likely to be an advantage rather than a limitation in this disease setting. Complete ablation of *VEGFA* in adult RPE causes choriocapillaris loss and cone dysfunction ^11^, and *VEGFA* blockade accelerates retinal ganglion cell death ^12^, indicating that a homeostatic *VEGFA* pool must be preserved for outer retinal integrity. In contrast efficacy does not require complete suppression. Previous studies have shown that AAV-delivered LbCpf1 produced 6.5% indels in mouse RPE yet reduced laser-induced CNV by 42%, comparable to aflibercept and without cone dysfunction on ERG ^43^. Similarly, AAV-SpCas9 achieving 22.5% indels reduced *VEGFA* protein by 26.6% and CNV area by 22-31% ^44^. The editing frequency reported here falls within this effective range. It also remains well below complete gene ablation, which in conditional knockout models produces choriocapillaris degeneration and cone dysfunction ^11^. Because activation is light-dependent, moderate editing could be further confined to the irradiated lesion area, leaving surrounding RPE unedited and able to sustain the *VEGFA* expression required by the underlying choriocapillaris.

However, several limitations should be acknowledged. The current experiments were conducted with an LED source at 0.15 mW/cm², corresponding to a fluence of 0.036 J/cm². This is approximately 1,400-fold lower than the 50 J/cm² used in clinical verteporfin photodynamic therapy ^45, 46^, and cellular toxicity under these conditions was well tolerated. Nevertheless, because verteporfin retains its photosensitising properties after treatment and can be activated by ambient light, the clearance profile of the nanoparticles will need to be characterised to define the period during which patients would require light protection. In addition, the *in vivo* efficacy assessment was limited to Day 8, shortly after the peak of vascular leakage, and therefore reflects the early treatment response. Longer-term studies will be needed to assess the durability of the therapeutic effect. Translation will also require optimisation of light delivery and nanoparticle dosing, since the illumination geometry and vitreous volume of the human eye differ substantially from those used here.

Taken together, these findings establish a foundation for spatial control over *VEGFA* disruption in the retina. Such precision matters because *VEGFA* is not a purely pathological mediator but a constitutive trophic factor required by the choriocapillaris and retinal function, and uncontrolled deletion removes that support irreversibly. By confining editing to the irradiated lesion, the approach could achieve localised suppression while leaving surrounding tissue intact. Precise genome editing with incomplete suppression may therefore overcome a key safety barrier to gene therapy in *VEGFA*-driven AMD and other neovascular macular diseases.

## Materials and Methods

### Materials

DOPE, DOTAP and cholesterol were ordered from Avanti Polar Lipids (Alabaster, USA). Verteporfin (VP; SML0534), and Pur-A-LyzerTM Maxi Dialysis Kit (Molecular weight cut-off 12-14 kDa; PURX12015) were ordered from Sigma-Aldrich (Sydney, Australia). Dulbecco’s Phosphate-Buffered Saline solution (DPBS, 14190250), Invitrogen™ Live Cell Imaging Solution (1X, A14291DJ), NucBlue™ Live Ready Probes™ Reagent (Hoechst 33342, R37605), AmpliTaq Gold 360 Master Mix (4398881), LIPOFECTAMINE CRISPRMAX 0.3ML EACH (Lipofectamine CRISPRMAX Cas9 Transfection Reagent) (CMAX00003), TRUECUT CAS9 PROTEIN V2,100UG 5UG/UL (TrueCut Cas9 Protein v2) (A36498), and sgRNA (A35534) were brought from Thermo Fisher Scientific (Australia). Human retinal pigment epithelial cell line (ARPE19, ATCCCL247) and DMEM: F-12 Medium (ATC302006) were purchased from the ATCC (Manassas, USA). Foetal Bovine Serum (FBS, SFBS) was purchased from Bovogen Biologicals Pty Ltd (Melbourne, Australia). DNeasy Blood & Tissue Kit (69504) was brought from Qiagen (Germany). CellTiter 96® AQueous One Solution Cell Proliferation Assay (MTS) kit (G3580) was ordered from Promega (WI, USA). Luna universal one-step TR-qPCR kit (E3005L), Monarch® PCR & DNA Cleanup Kit (5 μg) (T1030) and T7 endonuclease I (M0302S) were purchased from New England Biolabs (Ipswich, USA). Human VEGF Enzyme-Linked Immuno-sorbent Assay (ELISA) kits (ab222510) were brought from Abcam (Cambridge, MA). All primers were ordered from IDT (Coralville, USA). The gRNA sequences were listed in Table S1, primer sequences used for *VEGFA* RT-qPCR in ARPE19 cells were listed in Table S2, primer sequences used for *VEGFA* T7E1 assay were listed in Table S3, primers for NGS amplification were list in Table S4.

### Lipo-Cas9-VP preparation and characterisation

Lipo-Cas9-VP nanoparticles co-encapsulating Cas9/sgRNA and VP were formulated using a microfluidic mixing device. The organic phase was prepared by mixing DOTAP (25 mg/mL), DOPE (25 mg/mL), cholesterol (1 mg/mL), and VP (1 mg/mL), followed by dilution with ethanol to a final volume of 1 mL. The aqueous phase consisted of Cas9 (10 µM) and sgRNA002 (10 µM) diluted with Tris buffer. The aqueous and organic phases were combined using a microfluidic mixing device at a total flow rate of 12 mL/min and a 1:1 aqueous-to-organic flow rate ratio, resulting in Lipo-Cas9-VP nanoparticles. They were stored at 4°C until further use. Particle size distribution and zeta potential were measured using a Zetasizer Nano-ZS (Malvern Panalytical, Malvern, UK). The fluorescence properties of VP were characterised using a spectrofluorometer (FluoroMax-4, HORIBA Scientific, Kyoto, Japan). Nanoparticle morphology was examined by transmission electron microscopy (TEM; JEOL 1400, JEOL Ltd., Tokyo, Japan). Formulation stability was assessed by monitoring particle size and zeta potential over 12 days during storage at 4°C

### Assessment of cellular uptake

ARPE-19 human retinal pigment epithelial cells (1 × 10⁵ cells/mL) were seeded in glass-bottom petri dishes and incubated at 37°C for 24 h. After 24 h, the culture medium was removed, and the cells were incubated with 1 mL of Lipo-VP suspension for different incubation times. The cells were then washed twice with PBS (1×, pH 7.4) to remove unbound nanoparticles. Cellular uptake was imaged using a Leica SP2 confocal laser scanning microscope equipped with a 40× oil-immersion objective. A 405 nm violet laser was used to excite Hoechst 33342 and VP. Fluorescence emission was collected using a 405 nm channel for nuclear staining and a 680–710 nm band-pass filter for VP fluorescence. Cellular fluorescence was quantified using ImageJ software. The corrected total cell fluorescence (CTCF) was calculated using the following formula:

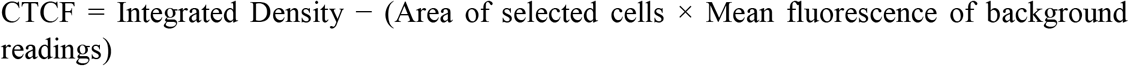

### Cellular viability assay under light illumination

The light power density was set at 0.15 mW/cm², with the light source positioned 1-3 cm from the ocular surface and maintained at a consistent distance throughout irradiation ARPE-19 cells were seeded in 96-well plates at a density of 2 × 10⁴ cells per well and cultured at 37°C for 24 h. The culture medium was then removed and replaced with fresh medium containing Lipo-VP at different concentrations. After 3 h of incubation, the Lipo-VP containing medium was removed, and the cells were washed once with fresh medium. The cells were then incubated with 100 µL of fresh medium and, where applicable, exposed to light illumination by LED (0.15 mW/cm²) at 690 nm for the indicated periods, followed by a further 24 h incubation. Cell viability was assessed using an MTS assay kit (Promega, WI, USA) according to the manufacturer’s instructions. After the 24 h incubation, the culture medium was removed, and 100 µL of fresh medium and 20 µL of MTS solution were added to each well. The cells were incubated for a further 1 h, after which the absorbance at 490 nm was measured using a microplate reader. Cell viability was expressed as a percentage relative to the untreated control and calculated using the following equation:

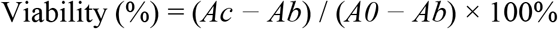

where *Ac* is the absorbance of the treated sample, *A0* is the absorbance of the untreated control, and *Ab* is the absorbance of the cell culture medium without cells.

### Stability study

Lipo-VP (100 µL) was diluted with 900 µL of cell culture medium (DMEM/F-12) containing either 0% or 10% FBS. For the positive control, 50 µL of Triton X-100 was added to disrupt the Lipo-VP. All samples were transferred into dialysis tubes (D-Tube Dialyzer Maxi, Merck KGaA, Germany). The dialysis devices were placed in 50 mL centrifuge tubes containing 13 mL of cell culture medium and maintained at room temperature under continuous gentle shaking. The fluorescence intensity of VP released from the nanoparticles into the surrounding medium was measured using a fluorescence spectrometer (FluoroMax, HORIBA Scientific, Japan) at 2, 6, 24 and 48 h. At 48 h, the total VP fluorescence was determined by disrupting the Lipo-VP with Triton X-100 and measuring the fluorescence after mixing the disrupted sample with the surrounding medium. The percentage of VP release was calculated using the following equation:

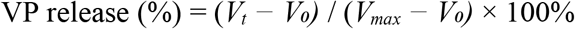

where *Vₜ* is the fluorescence intensity measured at each time point, *V₀* is the fluorescence intensity at 0 h, and *Vₘₐₓ* is the maximum fluorescence intensity measured after disruption of Lipo-VP with Triton X-100.

### RT-qPCR

*VEGFA* mRNA expression was quantified by RT-qPCR, with GAPDH used as the internal reference gene. Each sample was analysed in triplicate, using approximately 50 ng of total RNA per reaction. RT-qPCR was performed using the Luna Universal One-Step RT-qPCR Kit (E3005L, New England Biolabs) on a CFX96 Touch Real-Time PCR Detection System (Bio-Rad Laboratories, Inc.). The one-step RT-qPCR protocol consisted of reverse transcription at 55°C for 10 min, followed by an initial denaturation step at 95°C for 1 min and 40 amplification cycles of denaturation at 95°C for 10 s and annealing/extension at 60°C for 30 s. A melting curve analysis was performed to confirm the specificity of the amplified products. PCR products were further analysed by agarose gel electrophoresis to confirm the presence of a single band of the expected size. Relative *VEGFA* mRNA expression was calculated using the comparative quantification algorithms (2−ΔΔCt) and normalised to GAPDH expression, as previously described ^47^.

### PCR

The *VEGFA* genomic DNA sequence was analysed by PCR in triplicate. Genomic DNA extracted from ARPE-19 cells was used as the template. PCR was performed using AmpliTaq Gold 360 Master Mix (Thermo Fisher Scientific) on a CFX96 Touch Real-Time PCR Detection System (Bio-Rad Laboratories, Inc.). The PCR protocol consisted of an initial denaturation step at 95°C for 10 min, followed by 40 cycles of denaturation at 95°C for 30 s, annealing at 55°C for 30 s, and extension at 72°C for 30 s. A final extension was performed at 72°C for 7 min. The PCR products were purified using the Monarch PCR & DNA Cleanup Kit and analysed by agarose gel electrophoresis to confirm the presence of a single band of the expected size.

### T7 endonuclease I (T7E1) assay

A total of 200 ng of PCR product was denatured and re-annealed to form heteroduplexes using the following program: 95°C for 5 min, cooling from 95°C to 85°C at 2°C/s, followed by cooling from 85°C to 25°C at 0.1°C/s. The resulting heteroduplexes were digested with T7 endonuclease I (New England Biolabs) at 37°C for 15 min. The reaction was terminated by adding EDTA to a final concentration of 0.02 M. The digested products were separated by electrophoresis on a 2% TBE agarose gel. Band intensities were quantified using ImageJ software (National Institutes of Health, Bethesda, MD, USA). The percentage of indels was calculated using the following equation:

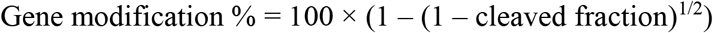

### ELISA assay

*VEGFA* protein levels were quantified using a Human VEGF ELISA Kit (Abcam, Cambridge, MA, USA) according to the manufacturer’s instructions. Treated cells were cultured in low-serum medium (DMEM/F-12 containing 1% FBS) for 3 h to minimise potential effects of serum on *VEGFA* measurements. The culture supernatants were then collected and centrifuged at 2,000 × *g* for 10 min to remove cellular debris. The clarified supernatants were carefully collected, avoiding the pellet, and used for ELISA analysis. A standard curve was generated by serial dilution of the *VEGFA* standards provided with the kit. Samples and standards were prepared using the appropriate diluents and added to the pre-coated microplate wells, followed by addition of the antibody cocktail. After incubation with shaking and subsequent washing steps, absorbance was measured using a SpectraMax iD5 microplate reader (Molecular Devices, USA). *VEGFA* concentrations were determined by interpolating the sample absorbance values from the standard curve.

### NGS

Genomic DNA was extracted from the treated ARPE-19 cells using the DNeasy Blood & Tissue Kit (Qiagen, Germany) according to the manufacturer’s instructions. A 358-bp genomic region spanning the target site was amplified using locus-specific primers containing Illumina Nextera overhang sequences (Table S4) and AmpliTaq Gold 360 Master Mix (Thermo Fisher Scientific, Australia). The resulting amplicons were supplied to the Ramaciotti Centre for Genomics (UNSW Sydney) for indexing PCR and paired-end sequencing (2 × 300 bp) on an Illumina MiSeq i100 Plus system. Sequencing reads were analysed using CRISPResso2 (version 2.3.4) and aligned to the 358-bp reference amplicon to quantify sequence modifications at the target site.

### Animal care and ethics

All animal procedures were reviewed and approved by the Animal Welfare and Ethics Committee (IACUC) of Saifu Laboratories Suzhou Co., Ltd (Ethics number: 2025-107-R for safety profile study and 2026-129-R for treatment efficiency study). Prior to study initiation, animals underwent an acclimation period of at least 5 days and were confirmed to be in suitable health for study inclusion. Animals were maintained in SPF facilities under controlled environmental conditions (20-26°C, 30-70% relative humidity, ≥15 air changes per hour, and a 12-h light/dark cycle). Animals were group-housed, with a maximum of three animals of the same sex and treatment group per cage, and provided SPF-grade rodent chow and purified drinking water ad libitum. Sterilized corn cob bedding was used. Animal health and welfare were monitored throughout the study by trained veterinary personnel, with appropriate veterinary care provided when required. Animal housing and care were conducted in accordance with GB14925-2023 and the recommendations of the Guide for the Care and Use of Laboratory Animals, 8th edition.

Specific pathogen-free (SPF) male Brown Norway (BN) rats were used in this study. BN rats were selected because they are widely used in experimental models of nAMD and exhibit well-characterised ocular phenotypes following CNV induction. Each animal was uniquely identified using an ear tag or microchip.

### Safety profile studies

Six male BN rats, aged 6–8 weeks, were randomly assigned to two groups to evaluate the ocular safety of Lipo-VP when administered by a single intravitreal injection to both eyes. The control group (n = 2) received PBS (vehicle; 2 µL/eye), while the treatment group (n = 4) received Lipo-VP (2 µL/eye; 0.5 µg/µL). The dosing day was defined as Day 1. Body weight was recorded before dosing, weekly throughout the dosing and recovery periods, and on the day of scheduled necropsy. Prior to ophthalmic examination, the pupils were dilated and animals were placed under general anaesthesia. Tropicamide eye drops (1-2 drops/eye) were administered for mydriasis. Anesthesia was induced via intramuscular injection of Zoletil® 50 (approximately 50 mg/kg) combined with xylazine hydrochloride (10 µL per animal). Ophthalmic examinations, including general ocular examination (GOE), intraocular pressure (IOP), fundus photography (FP), fundus fluorescence angiography (FFA), optical coherence tomography (OCT) and electroretinography (ERG), were performed once during the pre-dose phase for all animals, and once on Days 8 and 14, respectively. Histopathological examinations were conducted on Day 15 following euthanasia.

### Establishment of the laser-induced CNV model

CNV was induced by laser photocoagulation on Day 1 (D1) in both eyes of all animals. Rats were anesthetised and pupils were dilated before laser treatment. Laser photocoagulation was performed using a 532-nm green laser system (VISULAS 532s, Carl Zeiss, Germany) through a slit-lamp delivery system, with a handheld coverslip used as a contact lens to visualize the fundus. Four laser spots were applied around the optic nerve head of each eye, avoiding major retinal blood vessels. The laser parameters were set at 160 mW power, 100 μm spot diameter and 100 ms exposure duration.

Successful CNV induction was defined by the appearance of an intraoperative bubble at the laser site, indicating rupture of Bruch’s membrane, without evidence of preretinal or subretinal haemorrhage. Animals in which CNV induction was unsuccessful in one eye remained enrolled in the study, whereas animals with unsuccessful induction in both eyes were replaced with spare animals and the modelling procedure was repeated.

### Treatment efficacy evaluation

Eighteen male SPF BN rats, aged 6-8 weeks and weighing 140-300 g, with no significant abnormalities identified by clinical observation or ophthalmic examination, were enrolled in the study. Animals were randomly allocated into three groups, with body weight taken into consideration to achieve comparable distributions across groups: vehicle control, Lipo-Cas9-VP alone and Lipo-Cas9-VP + light illumination (n = 6 per group). On Day 1, laser-induced CNV was established in both eyes of all animals. Each animal received an intravitreal injection of 2 µL/eye of the assigned treatment. Approximately 30 min after completion of bilateral intravitreal injections, animals in the Lipo-Cas9-VP + light illumination group received 690-nm red-light irradiation. Both eyes were exposed with the eyelids naturally open. The light source positioned 1-3 cm from the ocular surface and maintained at a consistent distance throughout irradiation. Eyes were exposed to 690 nm light at an irradiance of 0.15 mW/cm² for 4 min, corresponding to a nominal light fluence of 0.036 J/cm². General clinical observations were performed daily throughout the study, and body weight was recorded weekly.

FP, FFA and OCT examinations were performed in all experimental animals on Days 1, 4 and 8. On Day 1, FP was performed immediately after laser-induced CNV modelling, while FFA and OCT assessments were performed on Days 4 and 8. Therapeutic efficacy was evaluated by monitoring fluorescein leakage on FFA and changes in the thickness of subretinal hyperreflective material (SHRM) on OCT over time. SHRM thickness at each laser-induced lesion was measured using the instrument’s built-in analysis software, and the mean SHRM thickness per eye was calculated. The primary efficacy endpoints were the mean fluorescein leakage score assessed by FFA and SHRM thickness measured by OCT.

### Statistical analysis

*In vitro* data are presented as the mean ± standard deviation (SD) or standard error of the mean (SEM), as indicated, from at least three independent experiments. Statistical analyses were performed using GraphPad Prism version 9.2.0 (GraphPad Software, San Diego, CA, USA). Comparisons between two groups were performed using two-tailed Student’s t-tests, whereas experiments involving multiple groups and/or experimental factors were analysed using two-way analysis of variance (ANOVA) followed by Tukey’s multiple-comparisons test, as appropriate.

*In vivo* quantitative data are presented as the mean ± SD or SEM, as indicated. For endpoints measured in both eyes, measurements from the left and right eyes were averaged for each rat to obtain a single animal-level value, with each rat considered one biological replicate. Qualitative ophthalmic assessments, including FP, FFA, OCT and histopathological findings, were summarised descriptively based on the incidence and/or severity of observed abnormalities, as appropriate.

For the ocular safety study, IOP and ERG measurements were compared between groups using unpaired Student’s t-tests or Mann-Whitney U tests, as appropriate. For the therapeutic efficacy study, longitudinal changes in quantitative endpoints were analysed using linear mixed-effects models, with treatment group, time and their interaction as fixed effects and animal as a random effect. Pairwise comparisons at each time point were adjusted using the Holm method. All tests were two-sided, with P ≤ 0.05 considered statistically significant. *In vivo* analyses were performed using Stata/IC version 15.1 (StataCorp LLC, College Station, TX, USA).

### Credit authorship contribution statement

**Rui Sang:** Data curation, Methodology, Formal analysis, Investigation, Writing - original draft, Writing - review & editing. **Xinpu Yang:** Data curation, Formal analysis, Investigation, Writing - review & editing. **Yagiz Alp Aksoy:** Conceptualization, Methodology, Formal analysis, Writing - review & editing. **Gerald Liew:** Methodology, Writing - review & editing. **Ewa M. Goldys:** Conceptualization, Resources, Funding acquisition, Supervision, Writing - review & editing. **Wei Deng:** Conceptualization, Methodology, Project administration, Resources, Funding acquisition, Supervision, Writing - review & editing.

### Declaration of interests

The authors report no conflicts of interest in this work.

## Acknowledgments

We acknowledge Electron Microscope Unit at Mark Wainwright Analytical Centre in UNSW for helping with the Cyro-TEM images and the Ramaciotti Centre for Genomics in UNSW for helping with the NGS. This work was financially supported by the project grant from Department of Education/Australia’s Economic Accelerator, Australia; UNSW Commercialisation Seed Fund and Australian Research Council Discovery Projects (DP240103024).

## Notes

### Competing Interest Statement

The authors have declared no competing interest.

