## Supplementary Information for "A light-responsive lipid nanoparticle platform delivering Cas9 RNP for *VEGFA* editing suppresses choroidal neovascularisation"

### **Method:**

#### **Optimisation and validation of *VEGFA* RT-qPCR primers**

Six candidate primer pairs targeting *VEGFA* were evaluated in human ARPE19 cells to identify an optimal primer set for quantitative gene-expression analysis. Total RNA (~100 ng per reaction) was analyzed using the Luna Universal One-Step RT-qPCR Kit (New England Biolabs) on a CFX96 Touch Real-Time PCR Detection System (Bio-Rad). Each 20-μL reaction contained 0.4 μM of each primers. Reverse transcription was performed at 55°C for 10 min, followed by initial denaturation at 95°C for 1 min and 40 cycles of 95°C for 10 s and 60°C for 30 s. *GAPDH* was used as the internal reference gene. Amplification profiles, melt-curve analysis and 3% agarose gel electrophoresis were used to assess amplification performance and product specificity. Candidate primer pairs with consistent amplification and predominantly specific products were further evaluated by Sanger sequencing to verify amplicon identity.

#### **PCR primer optimisation for the T7E1 assay**

Four candidate primer pairs targeting the genomic region encompassing the *VEGFA* CRISPR target site were evaluated using genomic DNA isolated from ARPE19 cells to identify an optimal primer set for the T7E1 assay. Approximately 100 ng of genomic DNA was used per reaction. PCR was performed using AmpliTaq Gold 360 Master Mix (Cat. No. 4398881, Thermo Fisher Scientific) on a CFX96 Real-Time PCR Detection System (Bio-Rad Laboratories, Inc.). The cycling conditions were 95°C for 10 min, followed by 40 cycles of 95°C for 30 s, 55°C for 30 s and 72°C for 30 s, with a final extension at 72°C for 7 min. PCR products were analysed by 3% agarose gel electrophoresis in TAE buffer at 100 V for 60 min to assess amplification and product specificity.

Table S1. gRNA sequences list

| Primer | Sequence 5'-3' |
| --- | --- |
| gRNA_hVEGFA001 | 5'- CCCAGCUACCACCUCCUCCC-3' |
| gRNA_hVEGFA002 | 5'- CGGGGAGGAGGUGGUAGCUG-3' |
| gRNA_hVEGFA003 | 5'- GCCGCCGGCCGGGGAGGAGG-3' |
| gRNA_hVEGFA004 | 5'- UCCGCCGCCGGCCGGGGAGG-3' |
| gRNA_VEGFA for rat study | 5'- GCCUCACCCGUCCAUGAGCC-3' |

Table S2. Primer sequences used for *VEGFA* RT-qPCR in human ARPE19 cells

| Primer | Sequence |
| --- | --- |
| hVEGFA_Seq1_F | 5'-GACGGACAGACAGACAGACA-3' |
| hVEGFA_Seq1_R | 5'-CGAGAACAGCCCAGAAGTTG-3' |
| hVEGFA_Seq2_F | 5'-GGACAGACAGACAGACACCG-3' |
| hVEGFA_Seq2_R | 5'-AAAAGTTTCAGTGCGACGCC-3' |
| hVEGFA_Seq3_F | 5'-GGAGGAAGAGTAGCTCGCCG-3' |
| hVEGFA_Seq3_R | 5'-GGAGGTAGAGCAGCAAGGCA-3' |
| hVEGFA_Seq4_F | 5'-CCCCAGCTACCACCTCCTC-3' |
| hVEGFA_Seq4_R | 5'-AAGCGAGAACAGCCCAGAAG-3' |
| hVEGFA_Seq5_F | 5'-GGAAGCCGAGCCGAGCGGAG-3' |
| hVEGFA_Seq5_R | 5'-GCGCGGCTGGAGCACTGTCT-3' |
| hVEGFA_Seq6_F | 5'-GACGGACAGACAGACAGACACC-3' |
| hVEGFA_Seq6_R | 5'-AAGCGAGAACAGCCCAGAAG-3' |
| GAPDH - F | 5'-CGCTCTCTGCTCCTCCTGTTC-3' |
| GAPDH - R | 5'-ATCCGTTGACTCCGACCTTCAC-3' |

Note: F: Forward primer R: Reverse primer.

Table S3. Primer sequences used for T7E1 assay

| Primer | Sequence |
| --- | --- |
| T7E1-Seq1_F | 5'-GACGGCTTGGGGAGATTGCT-3' |
| T7E1-Seq1_R | 5'-GCGAGAACAGCCCAGAAGTTG-3' |
| T7E1-Seq2_F | 5'-GCCATTCCCCACTTGAATCG-3' |
| T7E1-Seq2_R | 5'-CTCCTCCGAAGCGAGAACAG-3' |
| T7E1-Seq3_F | 5'-GGCTTGGGGAGATTGCTCTA-3' |
| T7E1-Seq3_R | 5'-CAGCCCAGAAGTTGGACGAA-3' |
| T7E1-Seq4_F | 5'-TGCTTGCCATTCCCCACTT-3' |
| T7E1-Seq4_R | 5'-AGCCCAGAAGTTGGACGAAA-3' |

Note: F: Forward primer R: Reverse primer.

Table S4. Primers for NGS amplification

| Primer | Sequence (5'-3') |
| --- | --- |
| *hVEGFANGS* - **Forward** | TCGTCGGCAGCGTCAGATGTGTATAAGAGACAG**AGA**<br>**GAGAAGTCGAGGAAGAGAG** |
| *hVEGFANGS* - **Reverse** | GTCTCGTGGGCTCGGAGATGTGTATAAGAGACAG**CT**<br>**CCTCCGAAGCGAGAACA** |

Illumina Nextera overhang sequences are shown in regular type; locus-specific sequences are shown in bold.

Table S5. Cas-OFFinder prediction analysis of potential off-target sites for the selected sgRNA sequence.

| Target | sgRNA sequence | Chromosome | Location | Predicted off-target sites |
| --- | --- | --- | --- | --- |
| *sgRNA_hVEGFA002*<br>(human source) | CGGGGAGGAGGUGGUAGCUG | Chr6 | 43770738 | 0 |
| *sgRNA_VEGFA*<br>(rat source for in vivo study) | GCCUCACCCGUCCAUGAGCC | Chr9 | 16233233 | 0 |

### Results:

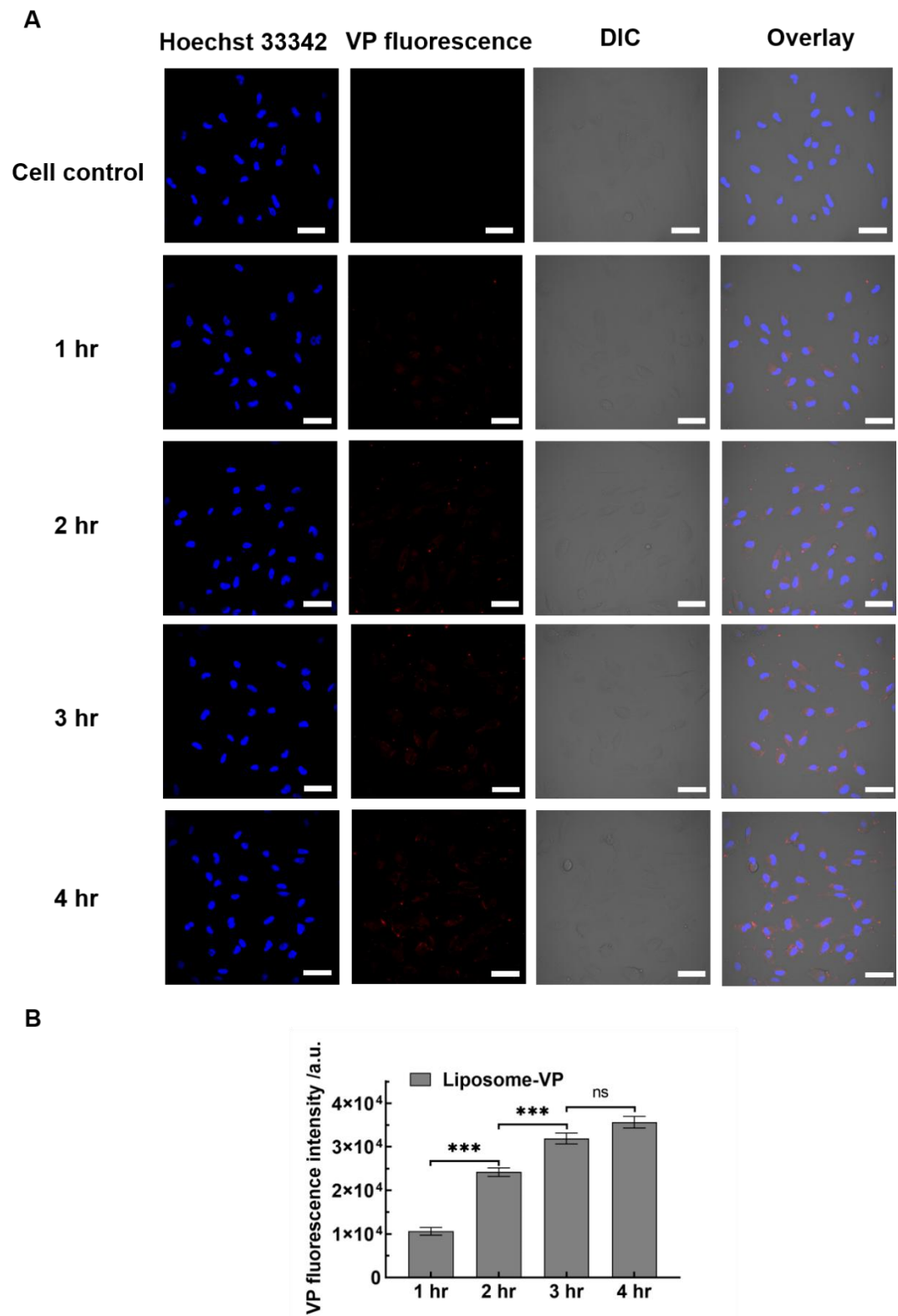

**Figure. S1** (A) Representative confocal microscope images of ARPE19 cells incubated with Lipo-VP (110.72 $\mu\text{g/mL}$) for 1 hr, 2 hr, 3hr and 4 hr. Scale bar is 50 $\mu\text{m}$. (B) Quantitative analysis of VP fluorescence from Lipo-VP indicated in Figure S1 (A).

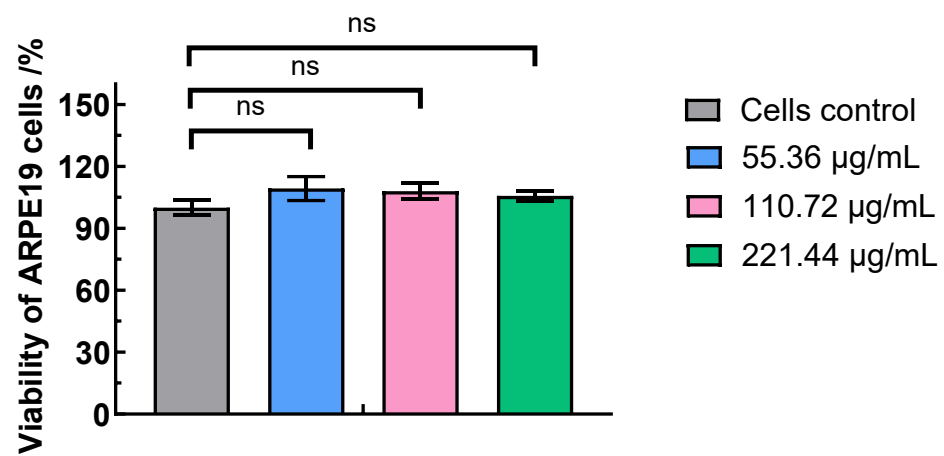

**Figure. S2** The viability percentage of ARPE19 cells at 24 hr after incubation with Lipo-VP for 3hr with different concentrations. (\*$p < 0.05$, \*\*$p < 0.01$ and \*\*\*$p < 0.005$; Mean ± SD, $n=3$).

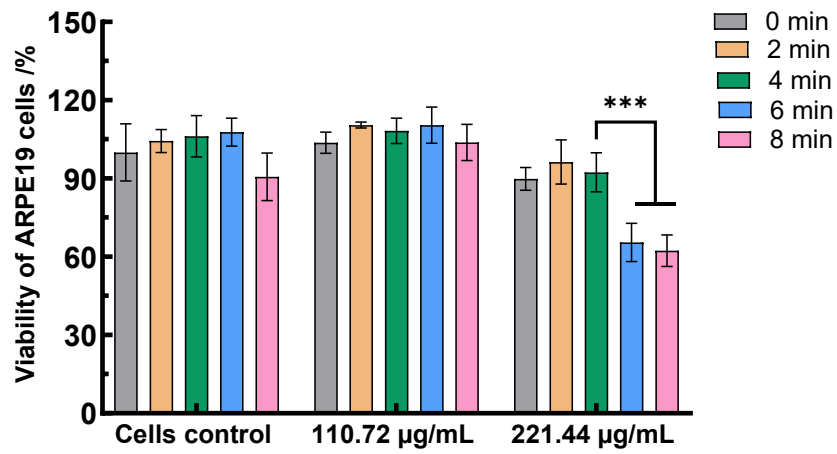

**Figure. S3** The viability percentage of ARPE19 cells at 24 hr after incubation with Lipo-VP at the indicated concentrations for 3 hr and exposed to light for 0-8 min. (\*$p < 0.05$, \*\*$p < 0.01$ and \*\*\*$p < 0.005$; Mean ± SD).

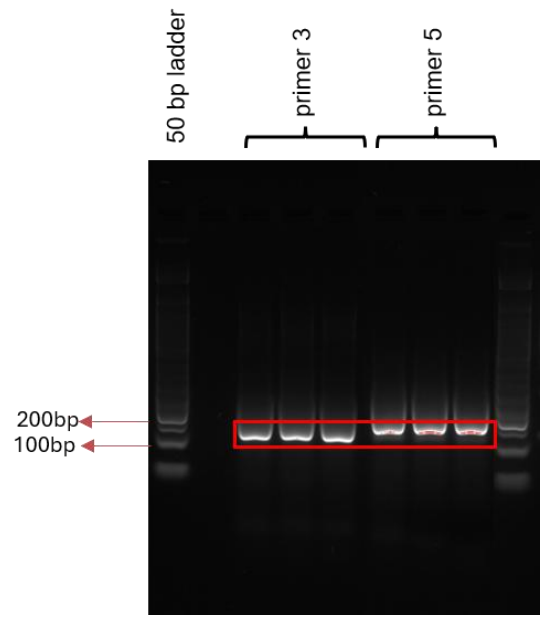

**Figure. S4** Agarose gel electrophoresis of RT-qPCR products for *VEGFA* expression in ARPE19 cells using primer pair 3 (161 bp) and primer pair 5 (186 bp). Products were separated on a 3% agarose gel at 100 V for 1 h.

Six candidate primer pairs targeting *VEGFA* were evaluated in ARPE19 cells for amplification performance and specificity. Comparison of amplification and melting profiles revealed differences in assay performance among the candidate primer pairs (Table S6). Amplicon size and specificity were further assessed by agarose gel electrophoresis (Figure S5). Primer pairs 3 and 5 showed consistent amplification and predominantly specific products and were selected for further sequence verification. Sanger sequencing confirmed the identity of the amplicon generated by primer pair 3, which showed high sequence identity to the intended *VEGFA* target. Based on the combined RT-qPCR, gel electrophoresis and sequence verification results, primer pair 3 was selected for subsequent *VEGFA* expression analysis.

**Table S6.** The melt temperature for 6 pairs of primer for RT-qPCR in ARPE19 cells

| Groups | VEGF<br>melt temperature<br>(°C) | GAPDH<br>melt temperature<br>(°C) |
| --- | --- | --- |
| Primer 1 | 76.50 | 85.00 |
|  | 88.00 | 85.00 |
|  | 81.50 | 85.00 |
| Primer 2 | 84.50 | 85.00 |
|  | 76.50 | 85.00 |
|  | 77.00 | 85.00 |
| Primer 3 | 92.50 | 85.00 |
|  | 92.50 | 85.00 |
|  | 92.50 | 85.00 |
| Primer 4 | 85.50 | 85.00 |
|  | 86.00 | 85.00 |
|  | 85.00 | 85.00 |
| Primer 5 | 90.50 | 85.00 |
|  | 90.50 | 85.00 |
|  | 90.50 | 85.00 |
| Primer 6 | None | 85.00 |
|  | None | 85.00 |
|  | 88.50 | 85.00 |

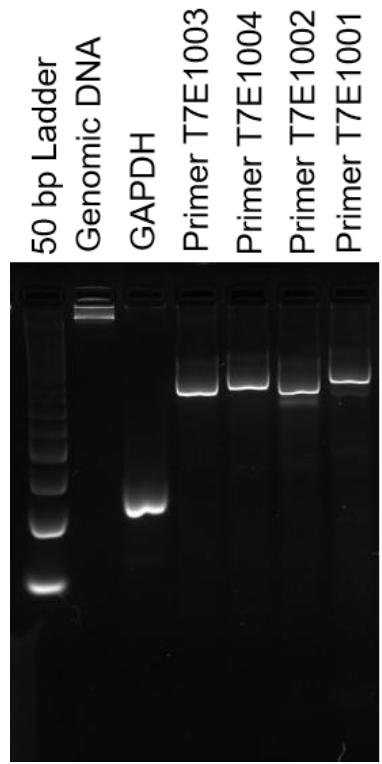

**Figure. S5** Agarose gel imaging of PCR products amplified by different primer pair of T7E1 assay for VEGF gene expression in ARPE19 cells, 3% agarose in TAE buffer, 100V 60 min.

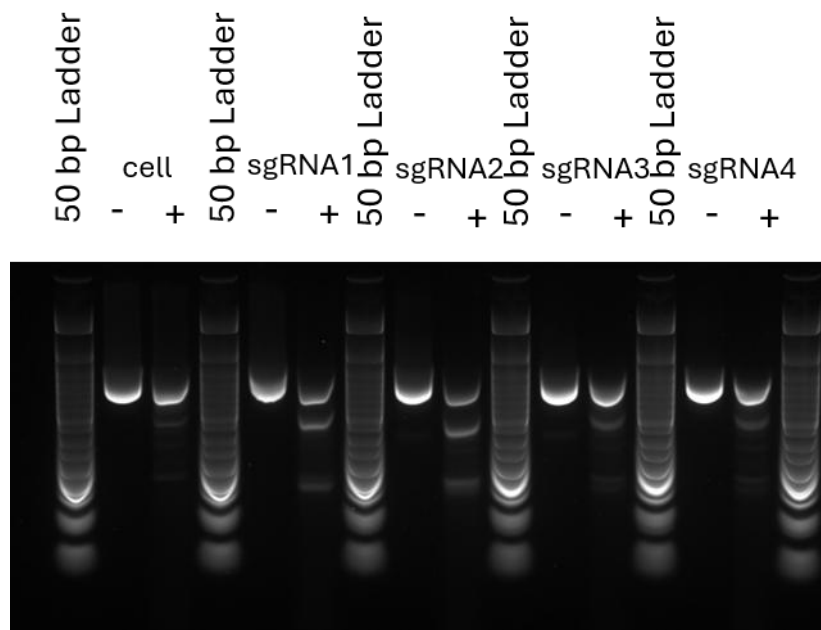

**Figure. S6** Agarose gel electrophoresis of PCR amplicons and T7E1 digestion products from ARPE19 cells transfected with Lipofectamine-CRISPR RNPs containing different gRNAs. Samples were collected 72 h post-transfection. Products were separated on a 2% agarose gel in TBE buffer at 100 V for 60 min.

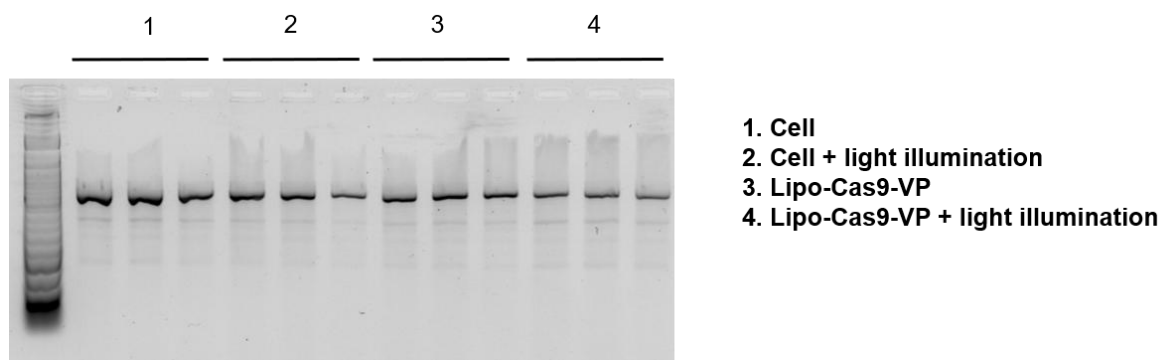

**Figure. S7** Agarose gel imaging of T7E1 products in ARPE19 cell lines under different treatments (2% agarose in TBE buffer, 100V 60 min).

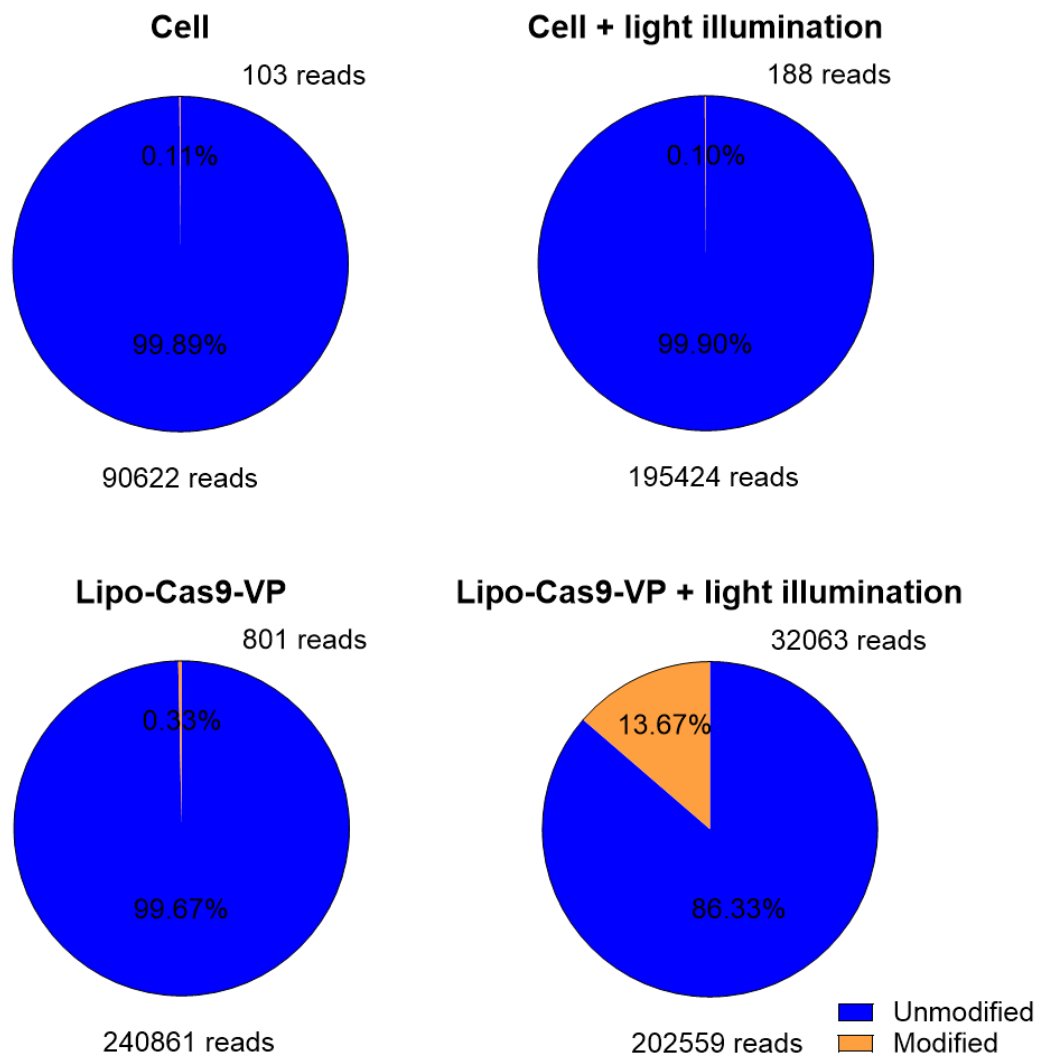

**Figure S8** NGS analysis of CRISPR-Cas9-mediated genome editing at the *VEGFA* target locus at 48 hr after the treatments. Pie charts show the proportions of unedited (blue) and edited (orange) reads in the indicated treatment groups. Editing frequencies were determined by sequence analysis using CRISPResso2.

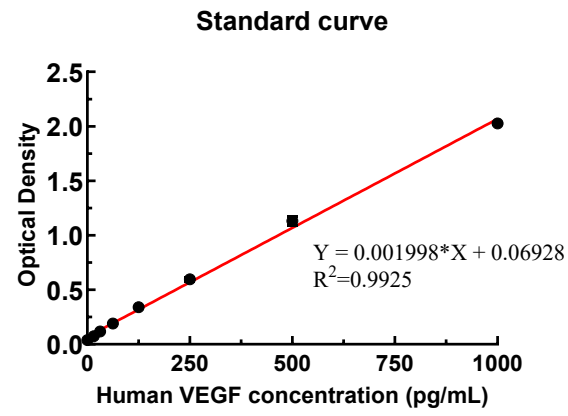

**Figure. S9** ELISA standard curve of VEGF protein. n=3.

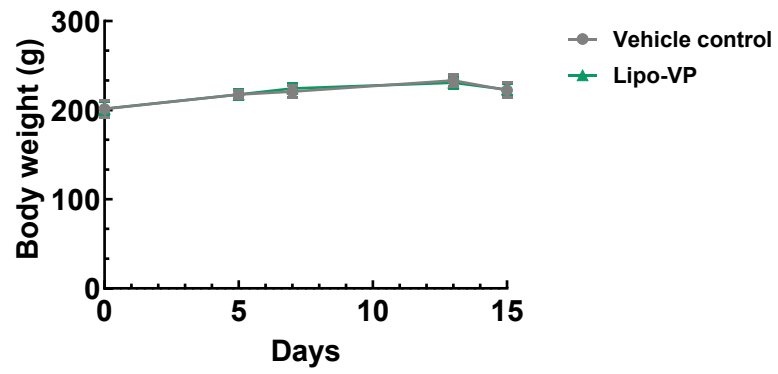

**Figure. S10** Body weight profiles of BN rats following intravitreal administration of Lipo-VP for *in vivo* ocular safety assessment.

**Table S7.** Summary of General Ophthalmic Examination

| Group | N (rats) | D0 | D8 | D14 |
| --- | --- | --- | --- | --- |
| Vehicle control | 2 | 0/2 abnormal | 0/2 abnormal | 0/2 abnormal |
| Lipo-VP | 4 | 0/4 abnormal | 0/4 abnormal | 0/4 abnormal |

**Table S8.** Summary of ERG Amplitude ($\mu\text{V}$) ($P \leq 0.05$)

| Days | Parameters | Vehicle Control N = 4 | Lipo-VP N = 8 |
| --- | --- | --- | --- |
| D0 | DA3.0-a wave | -121.65±31.11 | -137.48±47.24 |
|  | DA10.0-a wave | -127.98±20.93 | -137.15±32.34 |
|  | DA0.01-b wave | 153.95±27.58 | 178.28±31.12 |
|  | DA3.0-b wave | 507.88±104.28 | 490.54±109.6 |
|  | DA10.0-b wave | 537.48±65.02 | 566.98±91.62 |
| D8 | DA3.0-a wave | -56.10±32.22 | -91.38±33.24 |
|  | DA10.0-a wave | -55.93±38.21 | -100.31±33.81 |
|  | DA0.01-b wave | 52.05±25.00 | 97.06±38.23 |
|  | DA3.0-b wave | 185.38±68.31 | 329.93±103.00 |
|  | DA10.0-b wave | 204.43±87.62 | 385.41±101.82 |
| D14 | DA3.0-a wave | -166.10±31.84 | -72.88±28.62 |
|  | DA10.0-a wave | -147.88±25.16 | -99.13±28.34 |
|  | DA0.01-b wave | 119.63±14.49 | 91.50±20.70 |
|  | DA3.0-b wave | 487.85±59.41 | 293.53±41.61 |
|  | DA10.0-b wave | 505.58±36.67 | 381.31±78.09 |

**Peak Time:** During the experiment, no peak time changes of toxicological significance were observed. N = 4 eyes from 2 rats and N = 8 eyes from 4 rats.

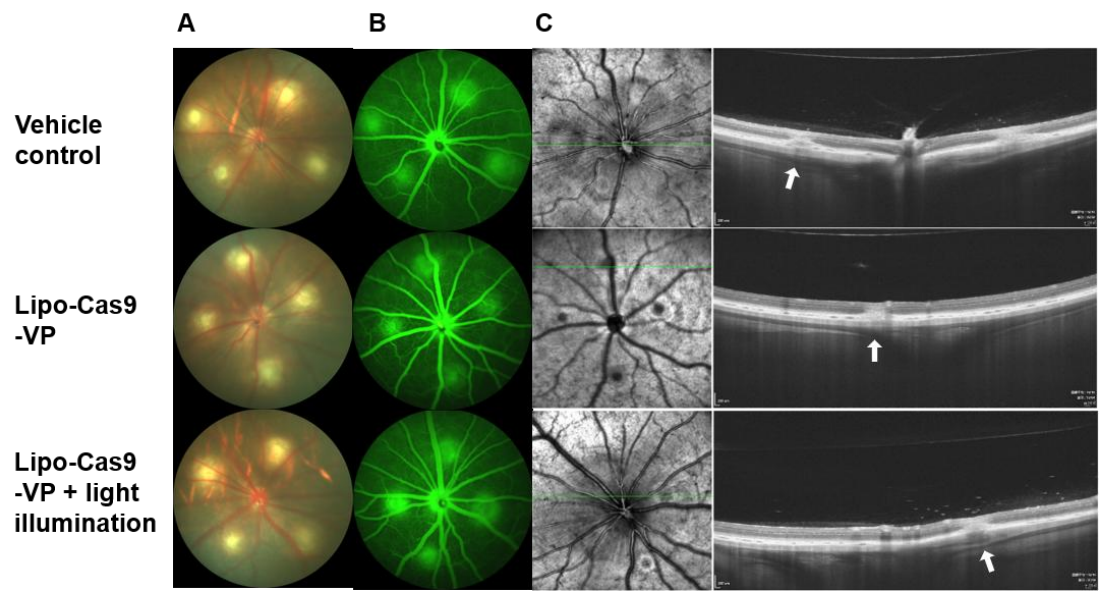

**Figure S11.** (A-C) Representative FP showing retinal appearance (A), FFA images (B) and OCT images (C) from the vehicle, Lipo-Cas9-VP and Lipo-Cas9-VP + light groups on day 4. The OCT scan through the optic disc was indicated by the green lines. White arrows indicate CNV-associated SHRM lesions on OCT. n = 6 rats.

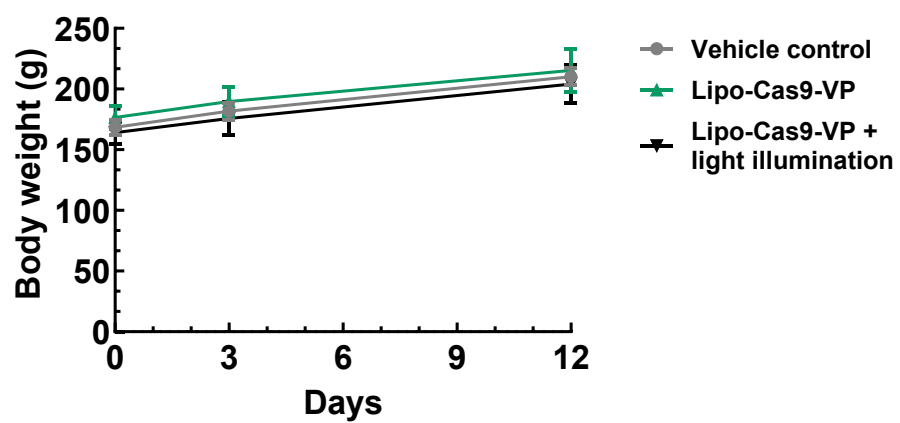

**Figure S12.** The body weight profile of BN rats following the indicated treatments. Data are presented as mean ± SD (N = 6 rats per group).
